# HNRNPH1-mediated splicing events regulate *EIF4G1* transcript variant composition and the organization of the *AURKA* 5’UTR

**DOI:** 10.1101/2025.07.28.667222

**Authors:** Tayvia Brownmiller, Soumya Sundara Rajan, Tamara L. Jones, Patricio Pichling, Vernon J. Ebegboni, Ashish Lal, Ioannis Grammatikakis, Erica C. Pehrsson, Natasha J. Caplen

**Affiliations:** Functional Genetics Section, Genetics Branch, Center for Cancer Research, NCI, NIH, Bethesda, MD 20892, USA; Omics Bioinformatics Facility, Genetics Branch, Center for Cancer Research, NCI, NIH, Bethesda, MD 20892, USA; Advanced Biomedical Computational Science, Frederick National Laboratory for Cancer Research, Frederick, MD 21702, USA; Regulatory RNAs and Cancer Section, Genetics Branch, Center for Cancer Research, NCI, NIH, Bethesda, MD 20892, USA

## Abstract

HNRNPH1 is a regulator of alternative splicing, but few studies have defined the splicing events it mediates. Here, we used short- and long-read RNA sequencing to interrogate the transcriptome-wide effects of HNRNPH1 depletion and its regulation of specific splicing events. Differential alternative splicing analysis revealed effects on the transcriptome that involved all splice event categories. We confirmed HNRNPH1’s regulation of a splicing event involving *TCF3*-exons 18a and 18b that encode distinct TCF3 transcription factor isoforms. Extending this finding, we present evidence that in neuroblastoma, *HNRNPH1* is a MYCN target, potentially explaining the higher levels of *HNRNPH1* and *TCF3-*exon 18a transcript variants in this tumor type. Analysis of two skipped exon events determined that HNRNPH1 regulates the splicing of exons encoding part of the EIF4G1 translation initiation factor’s N-terminus and an exon included in the 5’UTR of specific transcript variants encoding the mitotic kinase AURKA. Using reporter constructs, we show this *AURKA* 5’UTR exon enhances expression, suggesting HNRNPH1 could contribute to regulating AURKA protein levels. Our findings highlight HNRNPH1’s roles in regulating the expression of proteins with diverse cellular functions.

**Key points:**

- HNRNPH1 regulates the expression of proteins with diverse cellular functions, including proteins involved in the regulation of gene expression and essential cellular mechanisms.
- Depletion of HNRNPH1 alters the expression of specific protein-coding *EIF4G1* transcript variants.
- HNRNPH1 mediates the inclusion of an *AURKA* 5’UTR exon that enhances protein expression.

**Graphical Abstract:** 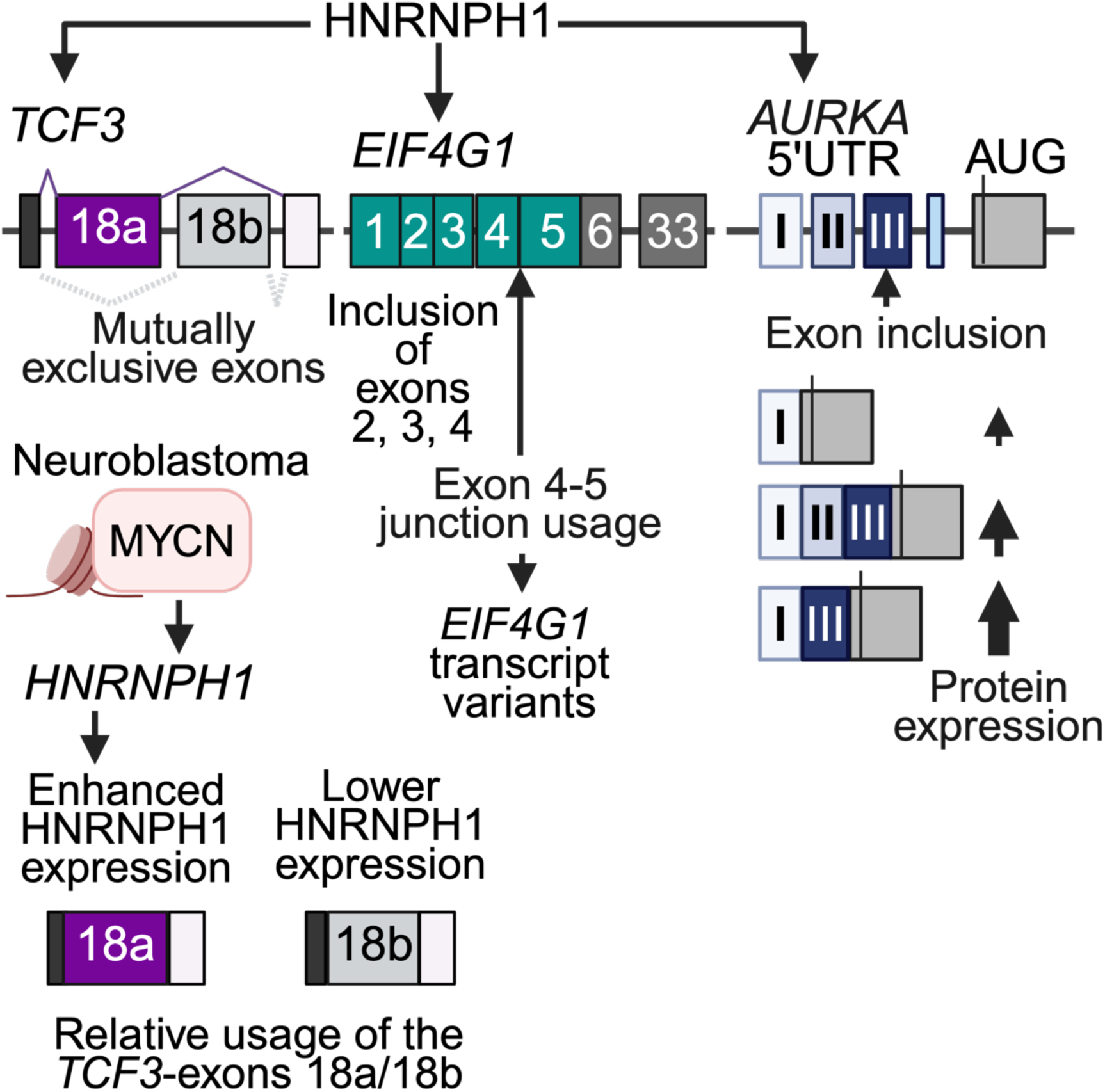

## Introduction

Splicing is a dynamic, highly regulated stepwise process that begins with assembly of the spliceosome on pre-mRNA via recognition of specific intronic sequences by the U1 and U2 small nuclear ribonucleoproteins (1,2). Working in concert with constitutive splicing, alternative splicing mechanisms result in the inclusion or exclusion of specific exons and variations in the incorporation or omission of other RNA sequences. Together, these processes underpin the diversity of expressed transcripts and thus the multiplicity of proteins needed for normal cellular function (3). The interaction of *tran*s-acting RNA-binding proteins (RBPs), most commonly members of the serine/arginine-rich splicing factor (SRSF) and heterogenous nuclear ribonuclear protein (hnRNP) family of proteins, can activate or repress spliceosome assembly through the binding of exonic or intronic *cis*-regulatory sequences (4). Collectively, over 90% of human multi-exon genes undergo alternative splicing (5). It is thus, unsurprising that many disease states exhibit alterations in the expression or function of RBPs and changes in the expression of specific transcript variants (6–8), including members of the HNRNPF/H family of proteins (reviewed in (9). For example, several rare neurodevelopmental disorders result from mutations in genes encoding members of the HNRNPF/H family of hnRNPs (10–14). Furthermore, studies have implicated members of the HNRNPF/H RBP family, particularly HNRNPH1, in the pathogenesis of multiple cancer types, including mantle cell lymphoma (15), Burkitt lymphoma (16), prostate cancer (17–19) and Ewing sarcoma (EWS) (20–22).

Quasi-RNA recognition motif (qRRM) domains that preferentially bind G-rich sequences are the defining feature of the HNRNPF/H family of RBPs. The most studied family members, HNRNPF, HNRNPH1, and HNRNPH2, are highly homologous. HNRNPH1 and HNRNPH2 proteins differ by only nine amino acids with >90% homology, and these two family members are ∼75% homologous to HNRNPF (reviewed in (9). The best characterized functions of HNRNPF and HNRNPH1/2 proteins relate to alternative splicing, though the outcome of an HNRNPF/H protein-mediated splicing event depends on several factors, including the location of the *cis*-regulatory sequences they bind and their interaction with other proteins. For example, the binding of sequences upstream of intronic 3’ splice sites or within an exon typically promotes exon inclusion while the binding of HNRNPF/H proteins to sequences downstream of intronic 5’ splice sites is usually associated with promoting exon exclusion (23). Though the HNRNPF, HNRNPH1, and HNRNPH2 proteins are highly homologous, their functions are not necessarily synonymous, and recent studies have begun to delineate alternative splicing event specificity related to each family member. For example, specific splicing events regulated by HNRNPH1 include transcripts expressing the fusion oncogene associated with Ewing sarcoma *EWSR1::FLI1* (20–22) and the gene encoding the transcription factor TCF3 (16).

In the case of *TCF3*, previous studies have reported HNRNPH1’s regulation of a *TCF3*-exon 18a/exon 18b mutually exclusive exon (MXE) splicing event via its binding of a G-rich region within exon 18b (24,25). This MXE splicing event has functional consequences as it results in the expression of two distinct TCF3 protein isoforms, E12 and E47, encoded by the exon 18a- and exon 18b- containing *TCF3*-transcript variants, respectively (26). The E12 and E47 isoforms of the TCF3 basic-helix-loop-helix (bHLH) transcription factor (also known as E2A) have different dimerization properties that alter their binding of regulatory E-box sequences, and the relative abundance of these isoforms regulates the differentiation of several stem or progenitor cell subtypes (24,27–30). For example, in human embryonic stem cells (hESCs), relatively high levels of HNRNPH1 favors the inclusion of *TCF3*-exon 18a and the expression of the E12 isoform, while lower levels of HNRNPH1 promotes the inclusion of *TCF3*-exon 18b, which results in the enhanced relative expression of the E47 isoform and a more balanced expression of both the E12 and E47 isoforms (16,24,25). In hESCs, changes in the relative expression of the TCF3 isoforms promotes cell differentiation, via E47’s suppression of E-cadherin expression (24). The TCF3 isoforms also contribute to B- and T-lymphocyte development. For example, the study of mouse models deficient for either E12 or E47 has shown that E47 is essential for early B-cell development, while both E12 and E47 function in regulating the transcription of genes encoding immunoglobulin light chains (27). A further demonstration of the importance of the relative expression of the TCF3 isoforms comes from the study of mutations in the *TCF3*-exon 18b HNRNPH-binding site observed in a subset of Burkitt lymphomas. Disruption of HNRNPH1’s binding of *TCF3*-exon 18b enhances the expression of *TCF3*-exon 18b containing transcript variants, which shifts the relative ratio of the two TCF3 protein isoforms, promoting tumorigenesis (16).

Recent studies have also connected HNRNPH1 to the alternative splicing of the proto-oncogene *HRAS* and the expression of the canonical p21 and alternative p19 HRAS isoforms, with the latter expressed due to the inclusion of *HRAS*-exon 5, which encodes a premature termination codon (19). In brief, Chen *et al*. reported that while both the HNRNPH1 and HNRNPF RBPs associate with *HRAS* pre-mRNAs, cell-based experiments and the positive correlation of the percentage inclusion of *HRAS*-exon 5 and *HNRNPH1* gene expression suggest that HNRNPH1 has a greater effect on stimulating the inclusion of *HRAS*-exon 5 than HNRNPF. This study also associated these findings to the observation that in many tumor types, including prostate, ovarian, and lung cancer, *HRAS*- exon 5 inclusion negatively correlates with a measure of MYC transcriptional activity, and that MYC activity also negatively correlates with *HNRNPH1* gene expression, but positively correlates with the expression of *HNRNPF.* In contrast, acute myeloid leukemia samples show a converse relationship, that is, a positive correlation of MYC activity with *HNRNPH1* gene expression, but a negative correlation with the expression of *HNRNPF*. Based on these and other findings, Chen *et al*. propose a model in which MYC activity determines the relative expression of HNRNPF and HNRNPH and that this influences the relative expression of *HRAS* transcript variants. Furthermore, based on ENCODE project data generated using the HepG2 liver cancer cell line, this study reported that loss of HNRNPH1 results in changes in over two thousand skipped exon (SE) events, though most of these remain uncharacterized. In this study, we have begun to address this deficit, by conducting a systematic analysis of HNRNPH1-dependent alternative splicing events.

Here, we report that HNRNPH1 regulates over twenty-five thousand splicing events in HEK-293T cells and over five thousand in HT-1080 cells. Our follow up analysis focused on specific splicing events mediated by HNRNPH1, including the *TCF3*-exon 18a/18b MXE event and SE events affecting the transcripts expressed by the *NUMB*, *RBM6*, *EIF4G1*, and *AURKA* genes. In the case of *AURKA*, we demonstrate that HNRNPH1 regulates the inclusion of a specific *AURKA* 5’UTR exon and the stability of *AURKA* transcripts. Collectively, these results demonstrate the importance of HNRNPH1 in the regulation of gene expression and suggest how its deregulated expression or function may contribute to different disease states.

## Materials and Methods

### Reagents

Reagents, including antibodies, primers, and siRNAs are detailed in **Supplementary Table S1**.

### Cell lines

The HEK-293T and HT-1080 cells were purchased from ATCC (Manassas, VA) and the neuroblastoma cell lines, SK-N-FI and IMR-32 were obtained from Dr. Javed Khan, Oncogenomics Section, Genetics Branch, CCR, NCI. HEK-293T cells were grown in DMEM (Invitrogen, Thermo Fisher Scientific, Waltham, MA), SK-N-FI cells in RPMI-1640 (Invitrogen, Thermo Fisher Scientific), and HT-1080 and IMR-32 cell lines in EMEM (ATCC). Media was supplemented with 10% FBS (Invitrogen, Thermo Fisher Scientific) and Plasmocin prophylactic (InvivoGen, San Diego, CA) at 37°C, 5% CO2. Cell line identity was confirmed by STR fingerprinting (ATCC, see **Data availability**), and cells were checked regularly for mycoplasma contamination (Lonza, Walkersville, MD). The HEK-293T cells were derived from human epithelial kidney cells, and HT-1080 cells from human epithelial cells obtained from the connective tissue of a patient with fibrosarcoma. The HT-1080 cells harbor a gain-of-function Q61K *NRAS* mutation and a switch-of-function R132C IDH1 mutation. SK-N-FI cells originate from bone-marrow metastasis of neuroblastoma and harbor *NF1* and *TP53* mutations. IMR-32 cells originate from an abdominal neuroblastoma mass and harbor amplification of the *MYCN* gene.

### Constructs

#### *AURKA* minigene

We synthesized a DNA cassette with 5′ SalI and 3′ SpeI restriction site overhangs (Genewiz, Azenta Life Sciences, South Plainfield, NJ). The cassette encompassed the full *AURKA* 5’UTR (including introns), coding exon 1, and 40 base pairs of intronic sequence upstream and downstream of these two features (**Supplemental Table S1)**. We cloned the DNA cassette into the pRHCglo splicing reporter plasmid (Addgene, Watertown, MA: 80169)(31). Oxford Nanopore Technologies (Oxford, UK) whole plasmid sequencing was used to confirm cloning and reporter cassette integrity.

### pNL2.2-5’UTR *AURKA* luciferase reporters

We synthesized DNA cassettes with 5′ NheI and 3′ HindIII restriction site overhangs (Genewiz, Azenta Life Sciences). These cassettes encompassed sequences corresponding to the *AURKA* 5’UTR exon combinations detailed in **Supplemental Table S1**. Each cassette was cloned into the pNL2.2[*NLucP*/Hygro] reporter plasmid (Promega), and we confirmed correct cloning using Sanger sequencing.

### RNAi and RNA analysis

To deplete HNRNPH1, we plated 250,000 cells per well of a six-well plate and transfected cells with 20 nM siRNA (siNeg (Qiagen, Germantown, MD), or siHNRNPH1 or siHNRNPH1(s6728), (Ambion, Thermo Fisher Scientific) complexed with 4.5 µL/well Lipofectamine RNAi-Max (Invitrogen, Thermo Fisher Scientific). We isolated RNA from three biological replicates 48-hours post-transfection using the Maxwell 16 LEV simplyRNA purification kit (Promega, Madison, WI) and gene silencing was confirmed using qPCR and the appropriate gene-specific primers detailed in **Supplementary Table S1**. Quantitative Real-Time PCR (qRT-PCR) was performed using PowerUp SYBR Green Master Mix (A25778; Thermo Fisher Scientific) on an ABI StepOne Plus Real-Time PCR system (Applied Biosystems, Foster City, CA). Fold-change gene expression was calculated by the ^ΔΔ^CT method and normalized to *NACA* or *RPL27* mRNA levels. For paired-end short-read RNA-seq, libraries were prepared from 1000 ng RNA (RIN > 9.0; three independent replicates) and sequenced using standard protocols and a Nextseq 2000 instrument (Illumina, San Diego, CA) as detailed previously (22). For long read RNA-seq (PacBio, Menlo Park, CA), we used 1000 ng of RNA pooled from three biological replicates (RIN > 9.0). cDNA library construction and sequencing were performed as previously described (22).

### RNA Stability Assays

Transfected cells were treated with either DMSO (D12345, Invitrogen, Thermo Fisher Scientific) or 5 µM of Actinomycin D (A1410, Sigma-Aldrich, St. Louis, MO; in DMSO), and harvested 0, 1.5, 3, 4.5, or 6 hours post treatment. RNA was harvested from control and treated cells and 1000 ng RNA used for cDNA synthesis. *AURKA* mRNA abundance was assessed using qRT-PCR designed to amplify all *AURKA* transcripts (*AURKA* exons 5-6). *GAPDH* was used for relative abundance normalization. Specific primer sequences are reported in **Supplementary Table S1.**

### Endogenous RNA splice assays

cDNA synthesis was performed using 1000 ng RNA and 5X iScript cDNA Supermix (BioRad, Hercules, CA). PCR amplification of *EIF4G1* and *TCF3* cDNAs were performed using the 2X Phusion High-Fidelity PCR Master Mix with GC Buffer (New England Biolabs, Ipswich, MA) and 2 µL of cDNA. PCR amplification of *AURKA* cDNAs was performed using 2X Q5 Hot Start High-Fidelity Master Mix (New England Biolabs) and 1 µL of cDNA. Specific cycling conditions are detailed in **Supplemental Table S1**. *EIF4G1* and *AURKA* amplified PCR products were resolved by electrophoresis using 6% TBE-PAGE gels (Invitrogen, Thermo Fisher Scientific). Sanger sequencing was used to confirm the sequences of amplified PCR products. *TCF3* PCR products were purified (Qiagen) and digested using the restriction enzyme PstI (New England Biolabs) before being resolved by electrophoresis using 6% TBE-PAGE gels. Gels were stained using SYBR safe DNA gel stain (Invitrogen, Thermo Fisher Scientific) and imaged on an Omega LumC (Aplegen Inc, Pleasanton. CA). Images were imported into ImageJ (32) for the quantification of each amplified product. The gel images shown are representative of three independent replicates and corresponding analysis and statistical analysis are indicative of these replicates.

### *AURKA* Minigene splicing analysis

We transfected HEK-293T cells (80,000 cells/well of a 12-well plate, grown overnight) with 0.8 μg of the *AURKA* 5’UTR minigene reporter (described above), with either no other nucleic acid or 20 nM siRNA using Lipofectamine 2000 (Invitrogen, Thermo Fisher Scientific). Transfected cells were grown for 48 hours and RNA extracted (Maxwell 16 LEV simplyRNA purification kit, Promega). First-strand cDNA synthesis was performed using 1000 ng RNA, the minigene specific primer (RTRHC - cDNA synthesis), and Superscript II RT (Invitrogen) reagents according to manufacturer’s instructions. For PCR amplification, we used the 2X Phusion High-Fidelity PCR Master Mix with GC Buffer (New England Biolabs), and 4-8 µL of the cDNA and 10 μM of each primer. **Supplemental Table S1** details the primer sequences and cycling conditions. PCR products were resolved by electrophoresis using 2% TBE agarose gels stained with SYBR safe DNA gel stain (Invitrogen, Thermo Fisher Scientific) and imaged on an Omega LumC (Aplegen). Images were imported into ImageJ for the quantification of each amplified product. The representative gel images and corresponding analysis shown are representative of three independent transfections.

### AURKA luciferase reporter assays

We co-transfected HEK-293T cells (10,000 cells/well; 96-well plate, grown overnight) with 60 ng of each pNL2.2-AURKA-5’UTR reporter (described above), 20 ng pGL4.54[*luc2*/TK that expresses Firefly luciferase (Promega), with either no other nucleic acid or 20 nM siRNA using Lipofectamine 2000 (Invitrogen, Thermo Fisher Scientific). After 48 hours, cells were processed using the Nano-Glo Dual Luciferase Reporter Assay System (Promega) according to the manufacturer’s 96-well plate, multi-channel pipette protocol. Luciferase signals (luminescence) were measured using a Perkin Elmer (Waltham, MA) Ensight plate reader.

### Immunoblotting

Cell pellets were collected and protein extracted using the Cell extraction buffer (Thermo Fisher Scientific: FNN0011) followed by brief sonication. Supernatants were collected and protein concentrations were assayed using BCA protein kit (Thermo Fisher Scientific: A55864). 20 - 30 µg of protein was loaded in each well and probed with indicated antibodies. Concentrations of the antibodies used are detailed in **Supplementary Table S1**. ImageJ was used for the quantification of the immunoblot intensities corresponding to specific protein isoforms.

### Computational analysis

Short-read RNA-seq data was processed using the CCR Collaborative Bioinformatics Resource (CCBR) RNAseq pipeline, RENEE (https://ccbr.github.io/RENEE/latest/#table-of-contents; https://zenodo.org/records/14502796). PacBio long read RNA-seq data was processed using the official Iso-Seq software, installed as a bioconda package. For each sample, CCS reads were processed using 1) Lima (parameters *--isoseq --peak-guess*) to remove primers and generate full-length reads, 2) IsoSeq refine (parameters *--require-polya*) to generate full-length non-chimeric reads, and 3) IsoSeq cluster (parameters *--use-qvs*) to cluster reads. High-quality clusters were mapped to the genome using the PacBio implementation of minimap2 (reference GRCh38 primary assembly, Gencode v41; parameters *--preset ISOSEQ --sort*), and redundant transcripts were collapsed into unique isoforms using IsoSeq collapse (parameters *--do-not-collapse-extra-5exons*). Pigeon was used to classify and filter the resulting isoforms (Gencode v41; classify parameter*-- gene-id*). For genes of interest, associated transcript sequences were extracted using the seqtk package (function *subseq*) and aligned to the reference genome using minimap2 (parameters *-ax splice -uf --secondary=no -C5*). Bam files were sorted and indexed using samtools. Processed long-read data was converted to MAJIQ-L compatible format in R using the package *stringr*. All alignments were performed using GRCh38.p13 v41 and transcript annotation performed using the corresponding Ensembl version (v107). Gene ontology analysis was performed using Metascape, (30). See **Supplementary Table S1** for further details of bioinformatic tools including version numbers.

### rMATS

Alternative splicing events were identified using rMATS (30). rMATS v1.4.2 was run with the parameters *-t paired --readLength 100 --libType fr-firststrand --novelSS --variable-read-length -- allow-clipping*. All relevant replicates were included in each comparison. rMATS JC.txt outputs were next filtered using the *maser* (<u>M</u>apping <u>A</u>lternative <u>S</u>plicing <u>E</u>vents to proteins) R/Bioconductor package (https://rdrr.io/bioc/maser/): 1) *filterByCoverage(data, avg_reads=5), 2) topEvents(data, fdr=0.05, delatPSI=0.1)*. Filtered rMATS event data were plotted using GraphPad Prism. Filtered rMATS events were visualized using the R package ggplot2 (volcano plots).

### Exon Enrichment

5’UTR, CDS, and 3’UTR coordinates were downloaded from the UCSC Table Brower (Gencode v41, comprehensive). rMATS target exons from significant SE and MXE events were intersected with the feature coordinates using bedtools. An exon was counted for each category if it overlapped any feature in that category from the same gene (minimum 1bp).

Enrichment was calculated as:

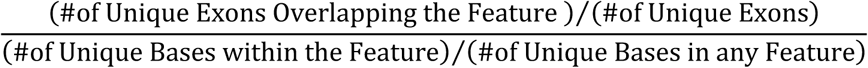

### MAJIQ, MAJIQ-L, and MAJIQlopedia

Local splicing variations (LSVs) were identified and quantified using the MAJIQ v2.5 and MAJIQ-L suite of tools (33,34). Splice-graphs were built using MAJIQ *build* with the parameter *--permissive*. Quantification for comparison across all short-read data was performed using the quantifier mode *heterogen* and run with two groups, siNeg and siHNRNPH1, with the parameter *--minreads* 5. Voila *tsv* was used to generate a “.tsv” readable file of the quantified LSVs for processing in R. For use with long-read RNA-seq data, the quantifier *psi* was implemented for each set of triplicate short-read sequencing experiments with the parameter *--minreads* 5. Voila *lr* was run using the matched psi and splice-graph output files from short-read data and IsoSeq long-read data to generate the long-read voila file using program defaults. Voila *view* was used to visualize all results. Splice-graph outputs were downloaded from the Voila *view*-generated browser in SVG format that were converted into Adobe Illustrator files. Normal and cancer summary data files were downloaded in “.tsv” format from the MAJIQlopedia splicing variation database (35). LSVs of interest were extracted using R and the data plotted using GraphPad Prism.

### Normalized Transcript Variant Percent Subset Abundance

Transcript variant counts for genes of interest were extracted from RSEM isoform counts using R. Excel was used for data analysis. Normalization was performed using a standard max-min approach, with substitution of the max value with the sum of transcript counts from protein coding transcript variants for a gene of interest.

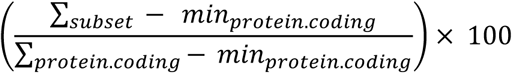

### Differential Transcript Variant Expression

A counts matrix for transcript variants was generated using RSEM isoform counts and used as input for the R package EBSeq (36). Analysis was performed using the EBSeq default settings. Results were annotated using Ensembl transcript information in R and data plotted using GraphPad Prism.

### Statistical Analyses

Calculations were performed in Excel (Microsoft), and data were exported to Prism 8.0.0 for Mac (GraphPad, La Jolla, CA) for statistical analyses. Unless stated otherwise, results are shown as mean ± standard error of the mean (SEM). A p-value of <0.05 was considered significant, though for some analyses, more stringent criteria were applied. Standard R functions and packages (ggplot2, tidyverse) were used to perform other analyses and to generate plots.

### External data sources

Neuroblastoma (NB) cell line transcriptome data (GSE89413) (37,38) were processed using the CCBR RNAseq pipeline, RENEE. For alternative splicing analysis, PSI quantification was performed using MAJIQ as described above. LSVs of interest were extracted using RStudio and associated data plotted using GraphPad Prism. NB cell line and tumor data supplied directly by Dr. Javed Khan, Genetics Branch, CCR, are described in Brohl *et al*. (38) (dbGAP phs001928 for pediatric cancers). Neuroblastoma cell line epigenome data (GSE138295 and GSE138314) was processed using the CCBR ChIPseq pipeline, CHAMPAGNE (https://ccbr.github.io/CHAMPAGNE/latest/), and loci of interest visualized with the Integrated Genome Viewer (IGV). Processed IMR-32 ChIPseq data (GSE184057) was downloaded and loci of interest visualized using IGV. Snapshots from IGV were exported and prepared for publication in Adobe Illustrator. SK-HEP-1 long-read RNA sequencing data (GSM8773037) was processed using IsoSeq as described above.

## Results

### The identification of HNRNPH1-regulated alternative splicing events

To identify high confidence splicing events regulated by HNRNPH1, we first conducted transcriptome-wide analysis of control and *HNRNPH1-*silenced HEK-293T and HT-1080 cells (**Supplementary Figure S1A**) using short-read RNA sequencing (RNA-seq) and employed the rMATS analytical tool (39) to determine significant changes in the following event types: skipped exon (SE), mutually exclusive exons (MXE), the use of an alternative 5’ (A5’SS) or 3’ splice site (A3’SS), or a retained intron (RI) (**Supplementary Table S2**). Quantification of the significantly affected alternative splicing events showed a greater number following the depletion of HNRNPH1 in HEK-293T cells than in HT-1080 cells (**Figure 1A**). We speculated that the higher expression of HNRNPH1 in HEK-293T cells than in HT-1080 cells (**Supplementary Figure S1B** and **S1C**) could, at least in part, explain this difference. To obtain supporting evidence for this hypothesis, we examined the documented *HRAS* HNRNPH1-dependent splicing event previously associated with the levels of HNRNPH1 expression, specifically the inclusion of *HRAS*-exon 5 that defines transcripts encoding the p19 HRAS isoform (19). Consistent with this previous study, depletion of HNRNPH1 in HEK-293T cells, which expresses relatively high levels of this RBP, resulted in decreased inclusion of *HRAS-*exon 5 (**Supplementary Figure S1D**). In contrast, HNRNPH1 depletion in HT-1080 cells, which express lower levels of HNRNPH1, relative to HEK- 293T cells showed no significant change in the inclusion of *HRAS-*exon 5. These data support the concept that the abundance of HNRNPH1 can contribute to its effects on alternative splicing, though we noted that HEK-293T and HT-1080 cells exhibited similar changes in the relative proportions of each alternative splicing event type (**Figure 1A**).

**Figure 1.**
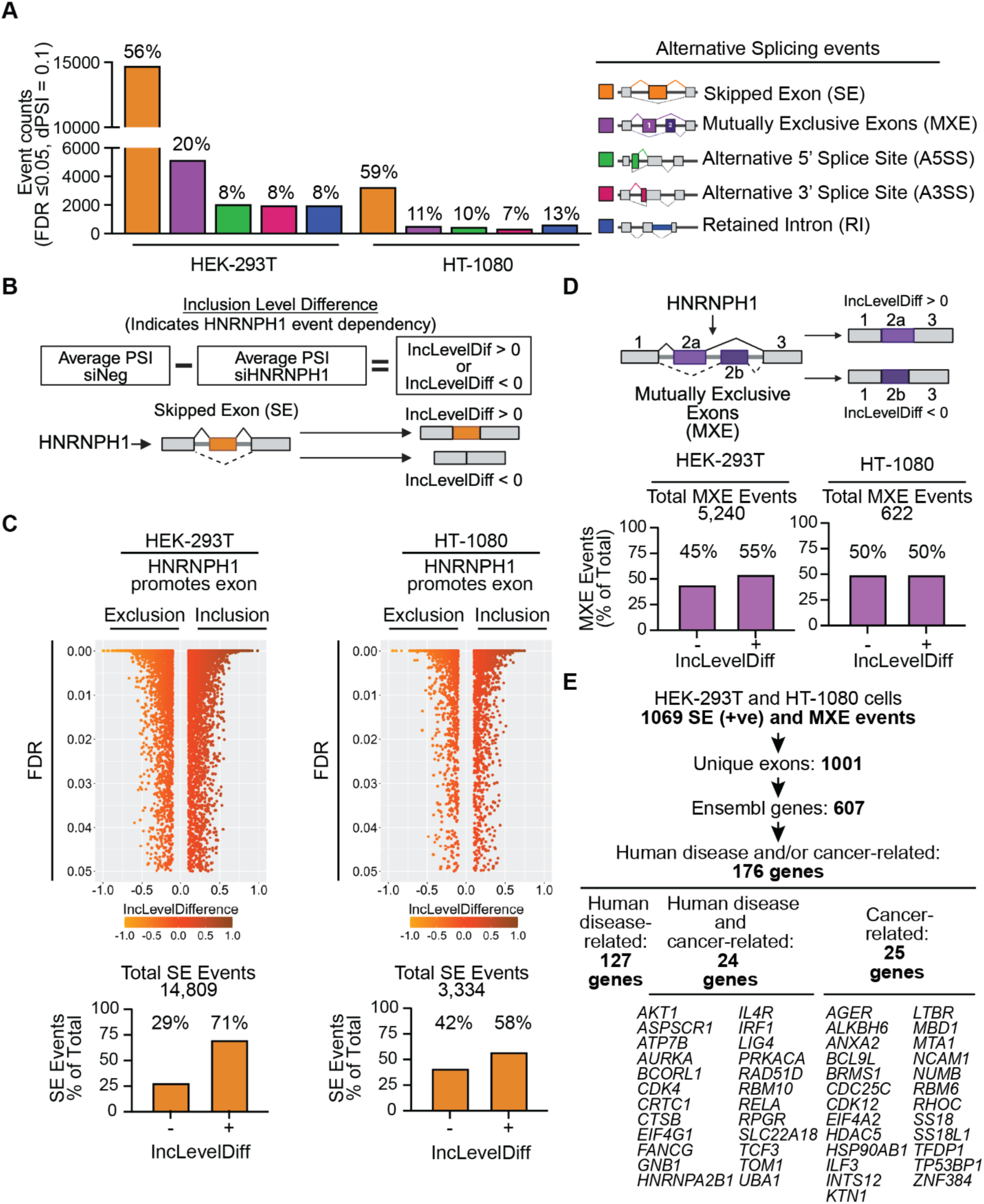
HNRNPH1’s transcriptome-wide regulation of alternative splicing. (**A**) The significantly affected alternative splice events following the silencing of *HNRNPH1* in HEK-293T or HT-1080 cells. (**B**) A schematic summarizing the analysis of inclusion level difference (IncLevelDiff) used to define HNRNPH1-dependent SE events. (**C**) Reverse volcano plots of the quantified SE events in each of the indicated cell lines, the total number of significant SE events detected (upper panel) and the percent distribution of IncLevelDiff values per cell line (lower panel). (**D**) A schematic summarizing the IncLevelDiff used to define HNRNPH1-dependent MXE splicing events (upper panel), the total number of significant MXE events (middle panel), and the percent distribution of IncLevelDiff values per cell line (lower panel). (**E**) Summary of the workflow used to prioritize genes (Ensembl gene symbol) for further analysis.

We next used the rMATS outputs from both cell lines as input for the R package *maser*. rMATS measures splicing changes by calculating the percent splice inclusion (PSI) and generates inclusion level difference (IncLevelDiff) values, and *maser* enables efficient handling of these data to interpret overall changes in alternative splicing events. The IncLevelDiff values obtained following the silencing of *HNRNPH1* indicates the predicted dependency of a given splice event on HNRNPH1 function. For example, in the context of SE events, an IncLevelDiff value of greater than zero (+) indicates that HNRNPH1 promotes the inclusion of an exon, while an IncLevelDiff value of less than zero (-) indicates that HNRNPH1 promotes the exclusion of an exon (**Figure 1B**). **Figure 1C** summarizes the distribution of the IncLevelDiff of all significantly affected SE events in the HEK-293T and HT-1080 cells following the silencing of *HNRNPH1*. We observed that ∼60-70% of the affected SE events have a positive IncLevelDiff value, demonstrating a bias towards HNRNPH1 promoting exon inclusion. Examination of the remaining four splice event types (**Figure 1D**, **Supplementary Figures S1E, S1F, S1G**) revealed, in most cases, similar positive and negative inclusion level differences across both cell lines, except for evidence that HNRNPH1 promotes greater use of the short form of an exon (A5’SS) in HT-1080 cells (**Supplementary Fig S1E**). Also, HNRNPH1-depletion in the HT-1080 cells resulted in detection of more RI (+) events compared to HEK-293T cells (**Supplementary Figure S1G**).

Skipped exon events with a positive IncLevelDiff, indicating HNRNPH1-dependent exon inclusion, represented the greatest proportion of high confidence splicing events, followed by MXE events, and we thus prioritized further analysis of these events. Using the workflow summarized in **Figure 1E**, we first defined the common MXE and positive IncLevelDiff SE events observed in both HEK- 293T and HT-1080 cells (**Supplementary Table S2**). This analysis defined 1069 shared events affecting 1001 unique exons, corresponding to 607 unique genes. These results suggest the presence of multiple splicing events regulated by HNRNPH1 in both cell line systems. Using the unique exons altered by one or more HNRNPH1-mediated splicing events, we next assessed whether these events showed enrichment for affecting the inclusion or exclusion of a protein-encoding exon versus an exon that contributes to untranslated regions (UTRs) (see Materials and Methods and **Supplementary Table S2**). Examining the 5’UTR, CDS, and 3’UTR gene regions as features, we observed a slightly greater enrichment over background for the altered usage of exons within protein-coding regions (SE: 2.16; MXE: 1.90), compared with exons that encode untranslated regions (5’UTR: SE: 1.26; MXE: 1.37; 3’UTR: SE: 1.40; MXE: 1.19). These results indicate that many HNRNPH1-regulated SE and/or MXE events have the potential to alter the identity and/or abundance of protein isoforms expressed by a particular gene. Gene ontology analysis of the 607 genes associated with an HNRNPH1-mediated SE or MXE event showed their enrichment for functions related to many different cellular processes, with the greatest enrichment for genes involved in the regulation of neurogenesis (**Supplementary Table S2, Supplementary Figure S1H**). Of these genes, this analysis highlighted 176 genes as functionally relevant to human disease and/or cancer or cancer-related mechanisms (**Figure 1E, Supplementary Table S2**), including *TCF3*.

### Transcriptome-wide analysis of HNRNPH1-dependent splicing events confirms its regulation of a *TCF3* MXE

Published studies (24,25) conducted using H9 human embryonic stem cells (hESCs) and Hela cells have shown HNRNPH1 regulates a *TCF3* MXE event involving the inclusion of either exon 18a (present in transcript variant *TCF3*-201 and three rarer variants) or exon 18b (present in transcript variant *TCF3*-211 and four rarer variants) (**Figure 2A, Supplementary Figure S2A**). Corroborating these studies, our rMATS analysis found that all the statistically significant *TCF3* splicing events altered following the depletion of HNRNPH1 in both HEK-293T and HT-1080 cells involved a reduction in the inclusion of *TCF3*-exon 18a (**Figure 2B and Supplementary Figure S2B**). To enable parallel evaluation of *TCF3*-exons 18a and 18b usage, we used a complementary suite of RNA-seq analysis tools called MAJIQ (33,40). MAJIQ uses a different approach than rMATS to quantify alternative splicing changes, accounting for the presence of more complex events that can occur within the same localized region by quantifying splice events based on all potential junctions to or from a reference exon. In the case of the *TCF3* MXE, the quantified outputs of MAJIQ, referred to as local splice variations or LSVs, enable the parallel assessment of changes in the inclusion of *TCF3-*exons 18a and 18b. Examination of the LSVs using *TCF3*-exon 17 as the reference exon, detected an increase in the inclusion of *TCF3*-exon 18b and a concomitant decrease in the inclusion of *TCF3*-exon 18a (**Figures 2C** and **2D, Supplementary Table S2**). To validate this finding, we used a PCR splice assay and confirmed that lower levels of HNRNPH1 favor the inclusion of *TCF3*-exon 18b over that of *TCF3*-exon 18a (**Figure 2E**). We observed similar results using a second siRNA targeting *HNRNPH1* (**Figure 2F and Supplementary Figure S2C**). Next, to assess whether the HNRNPH1-dependent regulation of splicing results in measurable changes in the expression of *TCF3* transcript variants, we employed two computational approaches: EBSeq, an analytical tool that identifies differentially expressed transcript variants in

**Figure 2:**
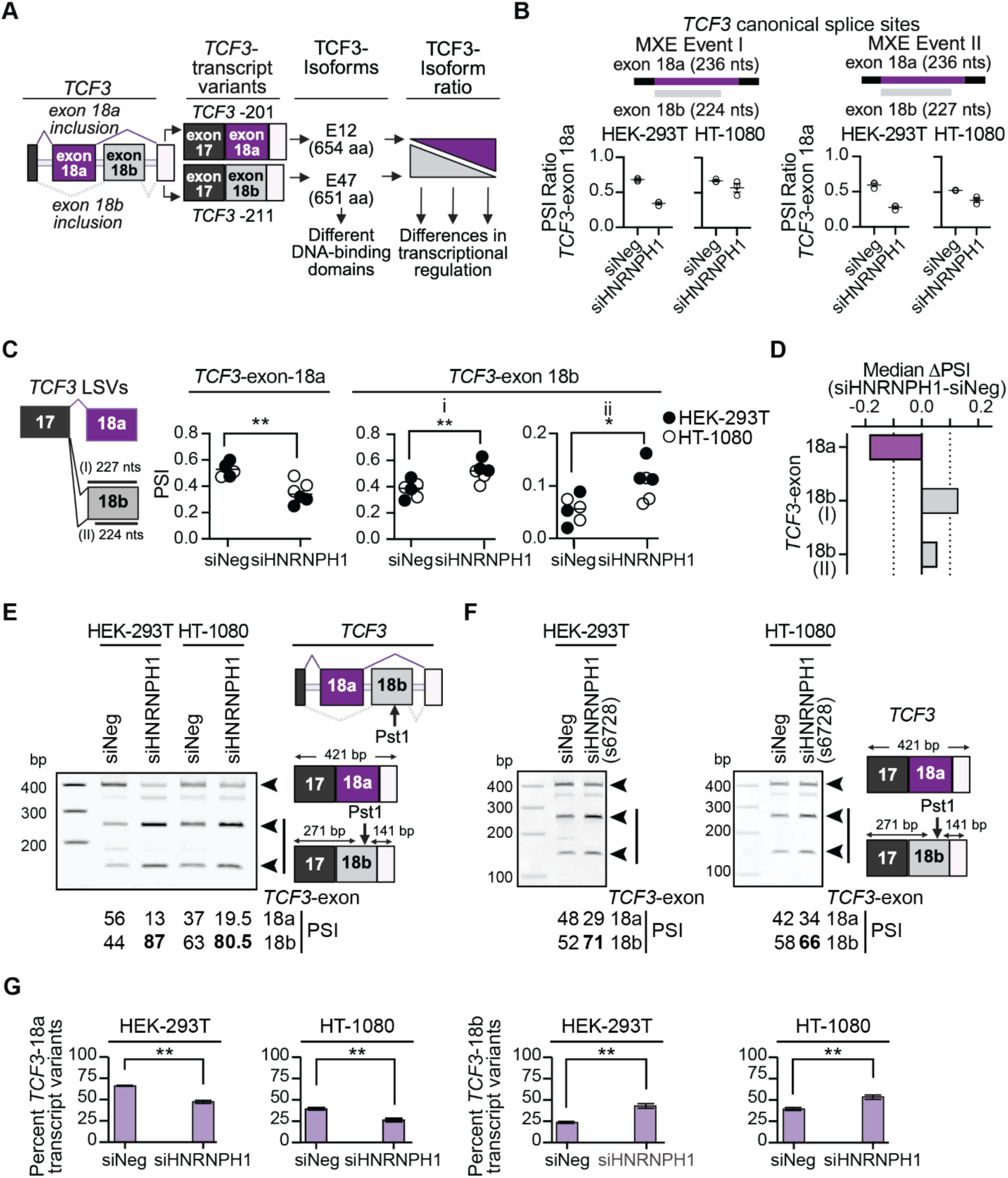
HNRNPH1’s regulation of a *TCF3* mutually exclusive splicing event. (**A**) Schematic of the *TCF3*-exon 18a and *TCF3*-exon 18b MXE, the canonical transcript variants defined by this MXE event, and their respective protein isoforms. (**B**) rMATS outputs for the *TCF3*-exon 18a and - exon 18b MXE splicing events that use canonical splice sites in the indicated cell lines transfected with the indicated siRNAs (three experimental replicas (open circles), mean and SEM (lines). The inclusion of the 224 nt or 227 nt versions of *TCF3*-exon 18b, respectively, define the MXE Events I and II. (**C**) Schematic of the relevant *TCF3*-exon 17, 18a, and 18b local splice variants (LSV; left hand panel) and the values generated using the heterogen (HET) quantifier within MAJIQ (33) (right hand panel). Statistical analysis: unpaired t test with Welch’s correction, P values * <0.05, ** <0.01). (**D**) Change in the inclusion of *TCF3*-exon 18a and *TCF3*-exon 18b (I) 18b-exon length 224 nts, (II) 18b-exon length 227 nts) following silencing of *HNRNPH1* quantified using the HET quantifier within MAJIQ. (**E**) PCR-based splicing assay (24) of the *TCF3*-exon 18a/18b MXE splicing event in control (siNeg) and HNRNPH1-depleted (siHNRNPH1) HEK-293T and HT-1080 cells. Data shown is representative of three independent experiments. (**F**) PCR-based splicing assay of the *TCF3*-exon 18a/18b MXE splicing event in control (siNeg) and HNRNPH1-depleted (siHNRNPH1(s6728)) HEK-293T and HT-1080 cells. Data shown is representative of three independent experiments. (**G**) Normalized percent subset abundance of *TCF3-*exon 18a (left) and -exon 18b (right) containing transcript variants expressed in control (siNeg) or HNRNPH1-depleted (siHNRNPH1) HEK-293T and HT-1080 cells based on RSEM counts (unpaired t test with Welch’s correction, P value ** <0.01).

RNA-seq experiments using empirical Bayesian statistical methods (36), (**Supplementary Figure S2D, Supplementary Table S2**) and normalized transcript variant percent subset abundance (see Materials and Methods) (**Figure 2G**). In both cell lines the overall abundance of *TCF3*-exon 18a containing transcripts decreased and *TCF3*-exon 18b containing transcripts increased following depletion of HNRNPH1 (**Figure 2G**), though the changes in the levels of individual transcript variants varied between them (**Supplementary Figure S2D**).

### Neuroblastomas exhibit relatively high levels of *HNRNPH1* and enhanced expression of *TCF3*-exon 18a containing transcript variants

Over 25% of Burkitt Lymphomas harbor mutations in *TCF3* (41,42). Interestingly, many of these mutations alter the HNRNPH-binding site present in *TCF3*-exon 18b, which results in promoting the inclusion of this exon over that of 18a, and thus, the expression of the TCF3 E47 isoform (16,42). There are no reports of other cancers harboring similar recurrent mutations, however, as recent studies have suggested that the levels of the HNRNPF/H proteins can contribute to tumorigenesis (19), we next examined whether the levels of *HNRNPH1* expression could promote alterations in the relative expression of *TCF3*-exon 18a or *TCF3*-exon 18b containing transcript variants across tumor types. To assess this hypothesis, we focused on data from solid tumors and used the recently published MAJIQlopedia resource that assesses LSVs across normal (GTEx) and tumor-derived transcriptome datasets (TCGA and TARGET) (35). Using this resource, we compared the relative ratio of *TCF3*-exon 18a/18b inclusion in normal and tumor tissues (**Figure 3A** and **Supplementary S3A**).

**Figure 3:**
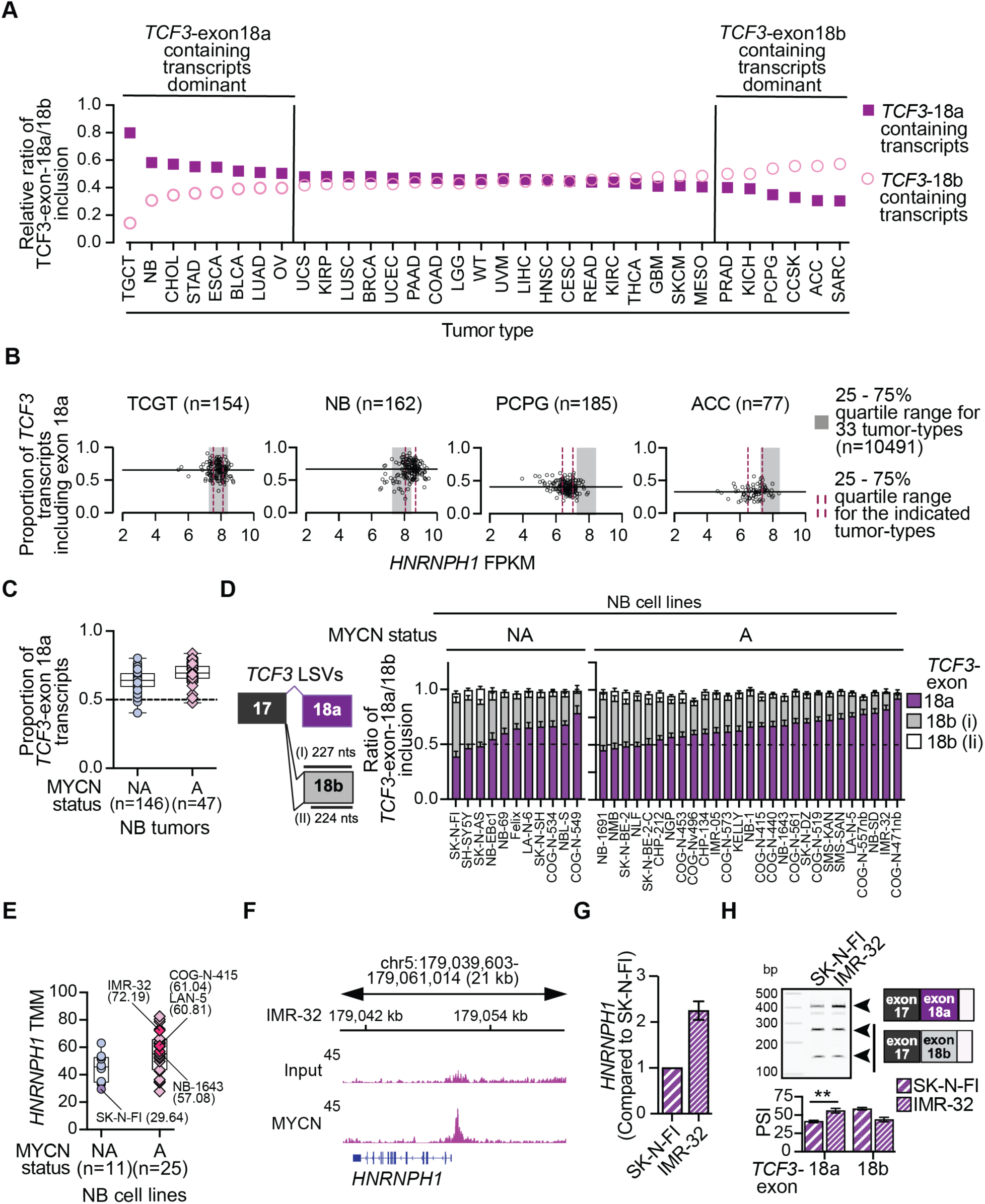
The differential expression of *TCF3* transcript variants in neuroblastoma. (**A**) Ranked relative ratios of *TCF3*-exon 18a and *TCF3*-exon 18b usage in the indicated tumor types. Data extracted from MAJIQlopedia. See **Supplementary Table S1** for the abbreviations used for each tumor type, the number of samples used for the analysis of each tumor type and the data source. (**B**) *HNRNPH1* gene level expression (all transcripts, FPKM) and the proportion of *TCF3* transcripts including *TCF3*-exon-18a for the indicated tumor types. The grey rectangle indicates the 25 – 75% quartile range of *HNRNPH1* expression in 10,491 samples representing 33 solid tumor types and the dotted red lines the 25 – 75% quartile range of *HNRNPH1* expression in the indicated tumor types. The black horizontal line indicates the median proportion of *TCF3* transcripts containing *TCF3*-exon-18a for the indicated tumor types. (**C**) The proportion of *TCF3* transcripts including exon 18a in NB tumors classified based on the indicated MYCN status. The black lines indicate the median and 25 – 75% quartile range. Data described in (38) and analyzed based on the expression of *TCF3* NM_003200 (*TCF3*-exon 18a) and *TCF3* NM_001136139 (*TCF3*-exon 18a). (**D**) The inclusion of exon 18a or exon 18b (I) exon length 224 nts; (II) exon length 227 nts) in *TCF3* transcripts expressed by the indicated NB cell lines. Data extracted from GSE89413 and analyzed using the PSI quantifier within MAJIQ. (**E**) The expression of *HNRNPH1* mRNA in NB cell lines categorized based on the non-amplification or amplification of MYCN (see (38) for additional details). Indicated are the names and *HNRNPH1* TMM values for NB cell lines mentioned in this study. (**F**) MYCN binding at the *HNRNPH1* locus of IMR-32 NB cells (Data extracted from GSE184057 (49). (**G**) qRT-PCR analysis of *HNRNPH1* mRNA expression in the IMR32 NB cell line normalized to that of SK-N-FI cells. (**H**) PCR-based splicing assay (24) of the *TCF3*-exon 18a/18b MXE splicing event using RNA prepared from SK-N-FI and IMR-32 NB cells. The image shown is representative of three RNA samples per cell lines and the graph shows the quantification (mean and SEM) of three samples per cell line.

Overall, most normal tissues expressed similar levels of *TCF3*-exon 18a and *TCF3*-exon 18b containing transcripts, though 17 tissue types showed evidence of greater *TCF3*-exon 18a inclusion (>10% difference), with bladder, skeletal muscle, naïve B-cells, and kidney cortex showing the greatest imbalance (**Supplementary Figure S3A**). Interestingly, several brain tissues showed a slight bias towards the inclusion of *TCF3*-exon 18b rather than *TCF3*-exon 18a, with those tissues that expressed more *HNRNPH1* exhibiting enhanced inclusion of *TCF3*-exon 18a, and those expressing less *HNRNPH1,* enhanced inclusion of *TCF3*-exon 18b (**Supplementary Figure S3B)**. Focusing next on 33 solid tumor types, most demonstrated similar levels of *TCF3*-exon 18a and exon 18b inclusion, but several tumor types exhibited a greater than 10% difference in the ratio of inclusion for these mutually exclusive exons (**Figure 3A**). For example, testicular germ cell tumors (TGCT) and neuroblastoma (NB) exhibit greater inclusion of *TCF3*-exon 18a than exon 18b, and pheochromocytoma and paragangliomas (PCPG) and adrenocortical carcinomas (ACC) exhibit the greater inclusion of *TCF3*-exon 18b than exon 18a.

Next, to assess whether the differential use of *TCF3*-exon 18a or exon 18b and their associated transcripts relates to the expression of *HNRNPH1* in TGCT, NB, PCPG, or ACC, we utilized the UCSC Toil RNAseq Recompute Compendium (hosted on Xena browser, 43) to reprocess TCGA and TARGET data using a standard pipeline starting with fastq files as the input. This analysis enabled the direct comparison of *HNRNPH1* gene level expression (all transcripts, FPKM) and the expression of exon 18a containing *TCF3* transcript variants (**Figure 3B**). First, to contextualize the expression of *HNRNPH1* in specific tumor types, we determined the range of its expression across cancers by calculating the 25-75% quartile range (indicated in **Figure 3B** as grey rectangles) using the data for the 33 solid tumor types we previously examined (>10,000 samples). Analysis of the TCGA and TARGET data confirmed the differential expression of *TCF3* transcript variants. Specifically, the TGCT and NB samples showed relatively high expression of *TCF3-*exon 18a containing transcripts, and PCPG and ACC showed relatively low expression of *TCF3-*exon 18a containing transcripts (median value indicated in each graph by the horizontal black line, **Figure 3B**). Examining the *HNRNPH1* gene-level data, the NB tumors expressed relatively high levels of *HNRNPH1* (25-75% quartile range indicated by red bars), while the TGCT samples express comparable levels of *HNRNPH1* to that seen across the >10,000 samples. In contrast, PCPG and ACC tumors expressed lower levels of *HNRNPH1*. The data generated from the NB, PCPG, and ACC samples are thus consistent with the previously described association of relatively high HNRNPH1 levels favoring the use of *TCF3-*exon 18a and lower levels resulting in the increased inclusion of *TCF3-*exon 18b (16,24,25). We next assessed the association of *HNRNPH1* abundance with *TCF3* transcript variant ratios in neuroblastoma in further detail.

Neuroblastoma derives from embryonic neural crest cells, and in NB tumors that harbor amplification of the proto-oncogene MYCN, a transcriptional core regulatory circuitry (CRC) that includes MYCN, HAND2, ISL1, PHOX2B, GATA3, and TBX2 promotes tumorigenesis (44). Biochemical studies have shown that the TCF3 E12 isoform can heterodimerize with the class II bHLH protein HAND2 (45) and the interaction of HAND2 and TCF3 (E2A) activates PHOX2B in the context of neuron differentiation (46). These studies suggest that the enhanced expression of *HNRNPH1* in NB relative to other tumor types and the dominant expression of *TCF3*-exon 18a transcripts observed in NB has the potential to influence the NB CRC. To assess this concept, we first examined *TCF3* transcript variant expression in NB tumors and cell lines (38,47) in the context of MYCN status (non-amplified or amplified). Examination of *TCF3* transcript variant expression showed that both MYCN non-amplified and amplified NB tumors express high levels of *TCF3*- exon 18a transcripts (as a proportion of all *TCF3* transcripts), with those tumors harboring the amplification of MYCN exhibiting greater expression than non-amplified tumors (**Figure 3C**). In addition, NB cell lines exhibited biased usage of *TCF3*-exon 18a over exon 18b (**Figure 3D**), and MYCN-amplified NB cell lines had a higher median expression of *HNRNPH1* than MYCN non-amplified cell lines (**Figure 3E**). To further interrogate a possible functional relationship between MYCN and *HNRPNH1* expression in NB, we examined deposited ChIP-Seq data from NB cell lines (48,49) and its binding at the *HNRNPH1* locus (**Figure 3F** and **Supplementary Figure S3C**). In all cases, these data showed evidence of MYCN binding at the *HNRNPH1* locus. Furthermore, analysis of data extracted from these same studies showed peaks corresponding to MYCN or MYC binding within sites of H3K4me3 modification that mark the transcriptional start sites of all the genes encoding the HNRNPF/H proteins (**Supplementary Table S3**). These findings support the recent transcriptome-based studies suggesting association of MYC activity and HNRNPF/H gene expression (19).

To confirm the differences in *HNRNPH1* expression exhibited by NB cells, we next used qRT-PCR to determine its expression in two NB cell lines: SK-N-FI, which is MYCN non-amplified and IMR-32 which is MYCN amplified. Consistent with the RNA-seq data, we observed that the IMR-32 cell line exhibits greater expression of *HNRNPH1* than SK-N-FI cells (**Figure 3G**). We observed similar differences at a protein level, though antibodies cannot distinguish HNRNPH1 and HNRNPH2 (**Supplementary Figure S3D**). We also assessed the relative usage of the *TCF3-*exon 18a and 18b using a PCR splice assay (**Figure 3H**) and observed enhanced inclusion of *TCF3*-exon 18a in IMR-32-derived RNA compared with RNA isolated from SK-N-FI cells. Collectively, these data suggest MYCN (or MYC) transcriptional activity in the context of NB has the potential to enhance the expression of HNRNPH1 which favors the expression of the E12 TCF3 isoform that interacts with at least one member of the NB CRC.

### The integration of short- and long-read transcriptome data identifies HNRNPH1-regulated splicing events that alter protein-coding transcripts

As our analysis of HNRNPH1’s regulation of the *TCF3* MXE splicing event validated our ability to detect HNRNPH1-dependent splicing events, we next prioritized analysis of uncharacterized HNRNPH1-dependent splicing events focusing on transcripts expressed by the human disease and/or cancer-related genes identified by our rMATS analysis (**Figure 1E**) and used our long-read RNA-seq data to identify those splicing events that affect full-length coding transcripts. In brief, we identified those genes for which long-read RNA-seq detected at least two transcript variants and total full-length read counts of greater than 200 under each condition, control (siNeg) and experimental (siHNRNPH1), and where the splicing event mapped to full-length transcripts with coding potential (**Supplementary Table S4**). This analysis identified splicing events affecting transcripts that included those encoding NUMB, EIF4G1, and AURKA.

The *NUMB* gene encodes a protein that functions as an inhibitor of Notch signaling and, via an interaction with MDM2, also functions as an indirect regulator of TP53 stability (reviewed in 50). Following the silencing of *HNRNPH1*, we observed significant skipping of *NUMB*-exon 12 (CDS exon 9). Specifically, integration of short- and long-read sequencing indicated a decrease in the use of *NUMB*-exon 11/12 and exon 12/13 junctions and increase in the use of *NUMB*-exon 11/13 junctions (**Supplementary Figure S4A**, black rectangles indicate key exon-exon junctions, and the arrows indicate the change in exon-exon junction usage following the silencing of *HNRNPH1*). Statistical analysis performed using rMATS (**Figure 4A**) and an assessment of LSVs by MAJIQ (**Figure 4B, Supplementary Table S4**) confirmed this observation, suggesting that HNRNPH1 participates in regulating the inclusion of *NUMB*-exon 12. This finding has functional consequences, as alternative splicing events define the generation of four major NUMB protein isoforms that differ in their functions. In the case of *NUMB*-exon 12, the transcript variants that include this exon encode the p72/71 isoforms, while the transcript variants encoding the p66/65 isoforms exclude this exon (51,52). The alternative splicing of *NUMB*-exon 12 contributes to whether the protein includes a 48 amino acid sequence within a disordered region that affects NUMB’s protein-protein interactions (reviewed in (50). Reflective of the importance of the *NUMB*- exon 12 splicing event, several studies have reported RBPs associated with its regulation, including RBM5 and RBM6 that promote its inclusion (53). Interestingly, we identified significant changes in the alternative splicing of *RBM6* transcripts following the depletion of HNRNPH1 (**Supplementary Table S4**) and our rMATS data revealed that HNRNPH1 depletion results in the decreased inclusion of *RBM6*-exon 6 (**Figure 4C**). Examination of the *RBM6* transcripts detected by long-read RNA-seq (see data availability to access long-read RNA-seq gff files) confirmed that depletion of HNRNPH1 alters the inclusion of *RBM6*-exon 6. Interestingly, *RBM6*-exon 6 encodes amino acids 495-519 of the canonical RBM6 protein, which includes a portion of an RRM domain (amino acids 456-536) (**Figure 4D**). We thus propose that depletion of HNRNPH1 will disrupt expression of fully functional RBM6. As previous studies have reported RBM6 as required for *NUMB-*exon 12 inclusion (53), we propose a model in which HNRNPH1 functions as an indirect regulator of the *NUMB-*exon 12 inclusion splicing event via RBM6 due to disruption of protein’s RRM domain and thus its interactions with RNA (**Figure 4E**).

**Figure 4:**
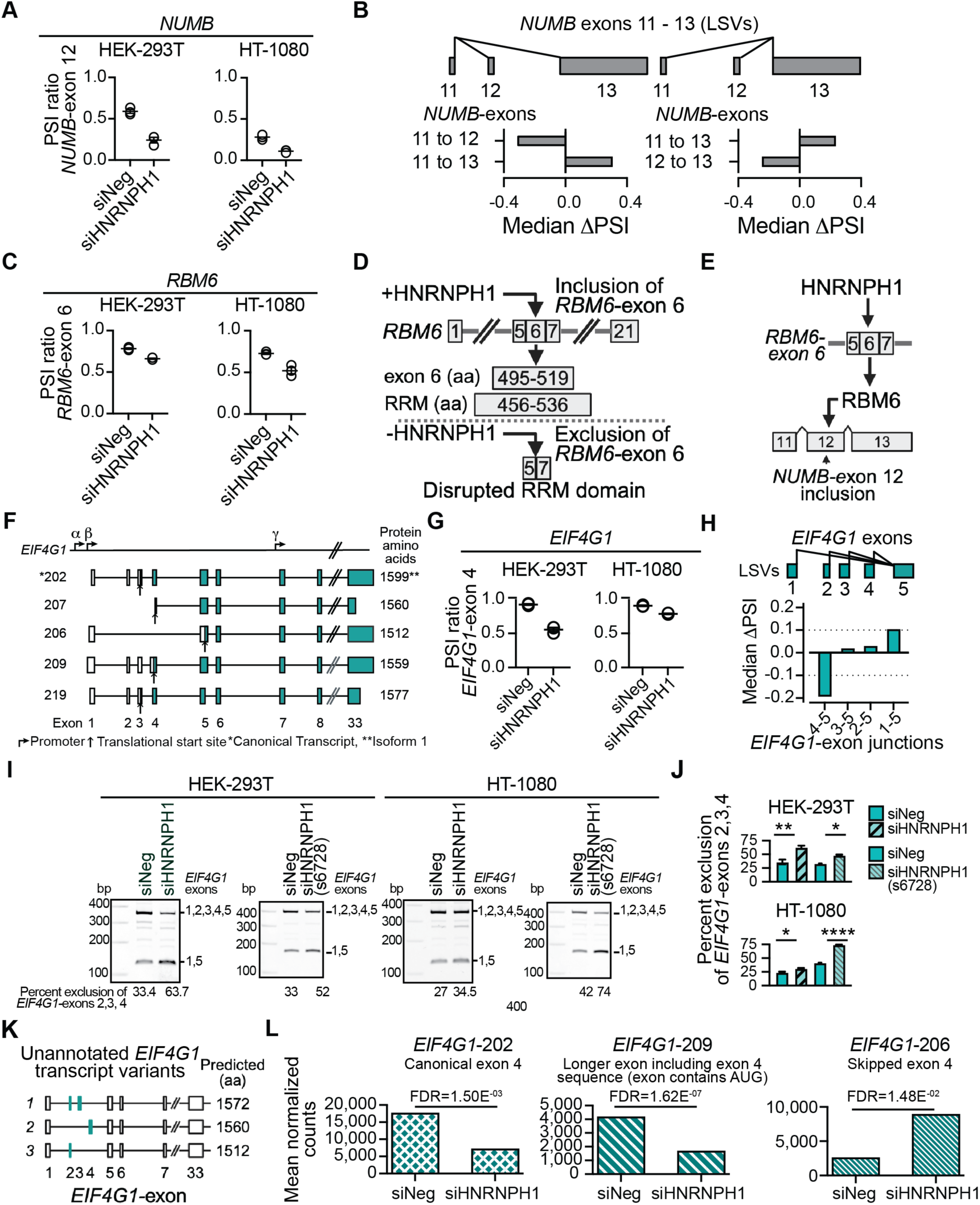
HNRNPH1-regulated splicing events affecting *NUMB*, *RBM6*, and *EIF4G1* transcripts. (**A**) rMATS outputs for the *NUMB*-exon 12 SE event in HEK-293T and HT-1080 cells transfected with the indicated siRNAs (three experimental replicas (open circles), mean and SEM (lines). (**B**) LSVs for *NUMB* exons 11-13 generated by the VOILA visualization tool and quantified using the HET quantifier within MAJIQ. (**C**) rMATS outputs for the *RBM6*-exon 6 SE event in HEK-293T cells transfected with the indicated siRNAs (three experimental replicas (open circles), mean and SEM (lines). (**D**) Model of HNRNPH1’s regulation of the *RBM6*-exon 6 SE splicing event. (**E**) A model of HNRNPH1’s proposed indirect regulation of the inclusion of *NUMB*-exon 12 via RBM6. (**F**) Schematic of selected *EIF4G1* transcripts focused on exons 1-8. The α, β, and γ indicate alternative transcriptional start sites., and the arrows below each transcript variant the translational start site. The white filled rectangles indicate untranslated regions, and the green filled rectangles, the protein-coding regions. (**G**) rMATS outputs for the *EIF4G1*-exon 4 SE event in the indicated cell lines transfected with the indicated siRNAs (three experimental replicas (open circles), mean and SEM (lines). (**H**) LSVs for the indicated *EIF4G1* exon-exon junctions generated by the VOILA visualization tool and quantified using the HET quantifier within MAJIQ. (**I**) Representative images of a PCR-based splice assay designed to detect the usage of *EIF4G1* exons 1 – 5 in HEK-293T or HT-1080 transfected cell lines (siNeg, siHNRNPH1 or siHNRNPH1(s6728)). The black lines specify the amplified products corresponding to the use of the indicated EIF4G1 exons. (**J**) Quantification of the *EIF4G1*-amplified product that excludes *EIF4G1* exons 2,3, and 4) in HEK-293T or HT-1080 transfected cell lines siNeg, siHNRNPH1, or siHNRNPH1(s6728)) determined using the *EIF4G1-* PCR-based splice assay (three independent transfections per condition per cell line, unpaired t test with Welch’s correction, P value * <0.05 ** <0.01, **** <0.0001. (**K**) Schematics of previously unreported full length *EIF4G1* transcripts focused on exons 1-10 based on long-read RNA-seq results from HEK-293T and HT-1080 cells and the predicted size of proteins encoded by these transcripts. (**L**) The mean normalized counts (EBSeq) of the indicated *EIF4G1* transcript variants in control (siNeg) and HNRNPH1-depleted (siHNRNPH1) HEK-293T.

The integration of our short and long-read RNA seq data also highlighted a putative HNRNPH1- dependent *EIF4G1* splicing event. EIF4G1 (eukaryotic translation initiation factor 4 gamma 1) functions as a scaffold for the formation of the EIF4F multi-subunit complex that recruits the 43S pre-initiation complex to the 5’ end of mRNAs to initiate protein translation (reviewed in 54). Currently, the 26 annotated *EIF4G1* transcript variants include 18 categorized as protein-coding, including those indicated in the schematic shown in **Figure 4F**. The regulation of EIF4G1 protein isoform expression includes the use of three different transcriptional promoters (α, β, and γ) and multiple translation initiation sites (55–57). A further determinant of these transcript variants involves the highly localized alternative splicing of *EIF4G1-*exons 1-9, yet the RBPs that facilitate these events remain poorly characterized. Consistent with previous studies, our results showed that the majority of *EIF4G1* transcripts originate from the α-promoter (see data availability to access long-read RNA-seq gff files) and exhibit alternative splicing centered on the first nine exons (**Supplementary Figure S4B**).

Following the silencing of *HNRNPH1*, we observed a decrease in the inclusion of *EIF4G1*-exon 4 (**Figure 4G** and **Supplementary Figure S4B**). MAJIQ analysis extended this finding by quantifying a decrease in the use of the *EIF4G1*-exon 4-5 junction and a concomitant increase in the usage of the *EIF4G1* 1-5, 2-5, or 3-5 exon-exon junctions, with the most significant increase involving the skipping of *EIF4G1*-exons 2, 3, and 4 (**Figure 4H, Supplementary Table S4**). To confirm this observation, we employed a PCR splice assay designed to detect transcript variants defined by the inclusion or exclusion of *EIF4G1-*exons 1 through 5 (**Figure 4I**). This assay detected dominant PCR products that correspond to amplification of cDNA containing *EIF4G1-*exons 1 through 5 (362 bp) and the skipping of *EIF4G1*-exons 2, 3, and 4 (124 bp). Consistent with our RNA-seq data, quantification of these dominant PCR products detected a decrease in the inclusion of *EIF4G1*- exons 2, 3, and 4 following HNRNPH1 depletion (**Figure 4J**).

Interestingly, the PCR assay also detected an amplified product corresponding to a rarer *EIF4G1* transcript variant (*EIF4G1-*226), and three other products of sizes corresponding to the usage of various combinations of the 5’ *EIF4G1* exons, specifically, exons 1, 2, 3, and 5, (* in **Supplementary Figure 4C**) 1, 3, and 5 or 1, 4, and 5 (** in **Supplementary Figure 4C**), and 1, 2, and 5 (*** in **Supplementary Figure 4C**). To assess if we had amplified mRNA corresponding to unannotated *EIF4G1* transcript variants, we examined our long-read RNA-seq data (see data availability to access long-read RNA-seq gff files and **Supplementary Table S4** for GenBank accessions) and identified full length reads corresponding to transcripts that use *EIF4G1*-exons 1, 2, and 3, *EIF4G1*- exons-1, 4, and 5, and *EIF4G1-*exons 1, 2, and 5 (**Figure 4K**). We further confirmed the detection of these transcripts in long-read RNA-seq data from a liver cancer cell line (SK-HEP-1) cells, see data availability to access long-read RNA-seq gff files and **Supplementary Table S4** for GenBank accessions). These unannotated *EIF4G1* transcript variants have the potential to encode protein isoforms that are slightly smaller than that of the canonical EIF4G1 protein (**Figure 4K**) and include predicted sequences matching UniProt database entries (unannotated variant 2: exons 2 and 3 excluded, UniProt Entry F74UU; unannotated variant 3: exons 3 and 4 excluded, UniProt Entry E9PGM1).

Both HEK-293T and HT-1080 cells showed decreased inclusion of *EIF4G1*-exon 4 following the silencing of *HNRNPH1* (**Figure 4G**). However, we focused on examining the effects of HNRNPH1- depletion on individual *EIF4G1* transcript variants using the data from HEK-293T cells (**Figure 4L**, **Supplementary Figure 4D**, and **Supplementary Table S4**) because these cells express much higher levels of *EIF4G1* than HT-1080 cells. The *EIF4G1*-exon 4 discussed above refers to annotations where this sequence forms part of the canonical *EIF4G1* transcript variant *EIF4G1*-202 and four other transcript variants, including *EIF4G1*-219. Five other *EIF4G1* transcript variants include the same sequence, but the use of alternative transcriptional start sites results in the annotation of this sequence as exon 2 in one transcript variant and exon 3 in four transcript variants. Four transcript variants, including *EIF4G1*-209 and *EIF4G1*-228, include the *EIF4G1*-exon 4 sequence as the 3’ end of a larger exon, and in each case, translation initiates within the exon 4 sequence. A further transcript variant, *EIF4G1*-207, includes the last 27 nts of *EIF4G1-*exon 4, with the first 3 nucleotides corresponding to the AUG codon. The depletion of HNRNPH1 in HEK-293T cells resulted in significant decreases in the abundance of the *EIF4G1*-202 and *EIF4G1*-209 transcript variants, but an increase in a transcript variant that skips *EIF4G1-*exons 2, 3, and 4 (*EIF4G1*-206) (**Figure 4L**). We observed similar changes in the abundance of other *EIF4G1* transcript variants, including those shown in **Supplementary Figure 4D**.

The shift to the exclusion of one or more exons upstream of *EIF4G1-*exon 5 is of interest because the N-terminus region of the human EIF4G1 protein (amino acids 1-200) contains the site required for EIF4G1’s interaction with the poly(A)tail-bound polyadenylate-binding protein (PABP) (58) (**Supplementary Figure S5A**). All the *EIF4G1* transcript variants highlighted by our analysis will encode an EIF4G1 protein that contains the minimal PAPB binding site. However, our data suggests that disrupted HNRNPH1 function, for example, because of mutation (12–14) could reduce the expression of multiple *EIF4G1* transcript variants, including the variant that encodes the full-length canonical EIF4G1 protein and the variants encoding isoforms expressed from a translational start site within exon 4. Such a shift in the composition of expressed *EIF4G1* transcript has the potential to promote the expression of transcript variants encoding an EIF4G1 isoform that lacks N-terminal amino acids present in the larger EIF4G1 proteins. The contribution of the N-terminal amino acids to EIF4G1 protein function is unclear. Nevertheless, we speculate that the proximity of the N-terminal portion of EIF4G1 to its PAPB region could impact interactions critical for the efficient initiation of translation and that any disruption of HNRNPH1’s function could thus have an indirect effect on translational efficiency (**Supplementary Figure 5B**).

### HNRNPH1 regulates the inclusion of an *AURKA* 5’UTR exon

We next examined the transcripts expressed by the *AURKA* gene, which encodes the essential serine/threonine kinase that regulates cell division. The integrated analysis of our short and long-read RNA-seq data (siNeg and siHNRNPH1) confirmed that the alternative splicing of *AURKA* transcripts centers on exons that form the 5’UTR of *AURKA* mRNAs (**Figure 5A** and see data availability to access long-read RNA-seq gff files). According to current annotation resources, four exons can contribute to the 5’UTR of *AURKA* transcripts, which, for convenience, we refer to here as 5’UTR exons I – IV (**Figure 5B**). A 395-nt genomic region encodes the 5’UTR exon I sequences incorporated into the annotated *AURKA* transcripts, but the size of this exon can vary between 29 and 279 nts in length depending on the transcript variant, with the most frequently used (and canonical) exon size being 48 nts. The most abundant *AURKA* transcripts, *AURKA-*206 and -208, along with *AURKA*-202 and -204, exclude any intervening exons, while other transcript variants (*AURKA*-201, -205, -207, -209, -210, -211, -212) include and exclude different combinations of 5’UTR exons II, III, or IV (**Figure 5B**). Seven transcript variants (*AURKA*-201, -202, -203, -205, -206, -207, -208) code for the same 403 amino acid protein, while the remaining six encode isoforms of 79 to 347 amino acids in length, with five encoded by mRNAs that use proximal polyadenylation signals and the sixth, the use of an alternative 3’UTR. Importantly, regulators of these alternative splicing events are unknown, and it is unclear how the expression of different transcript variants affects AURKA protein expression overall or the expression of specific AURKA isoforms.

**Figure 5:**
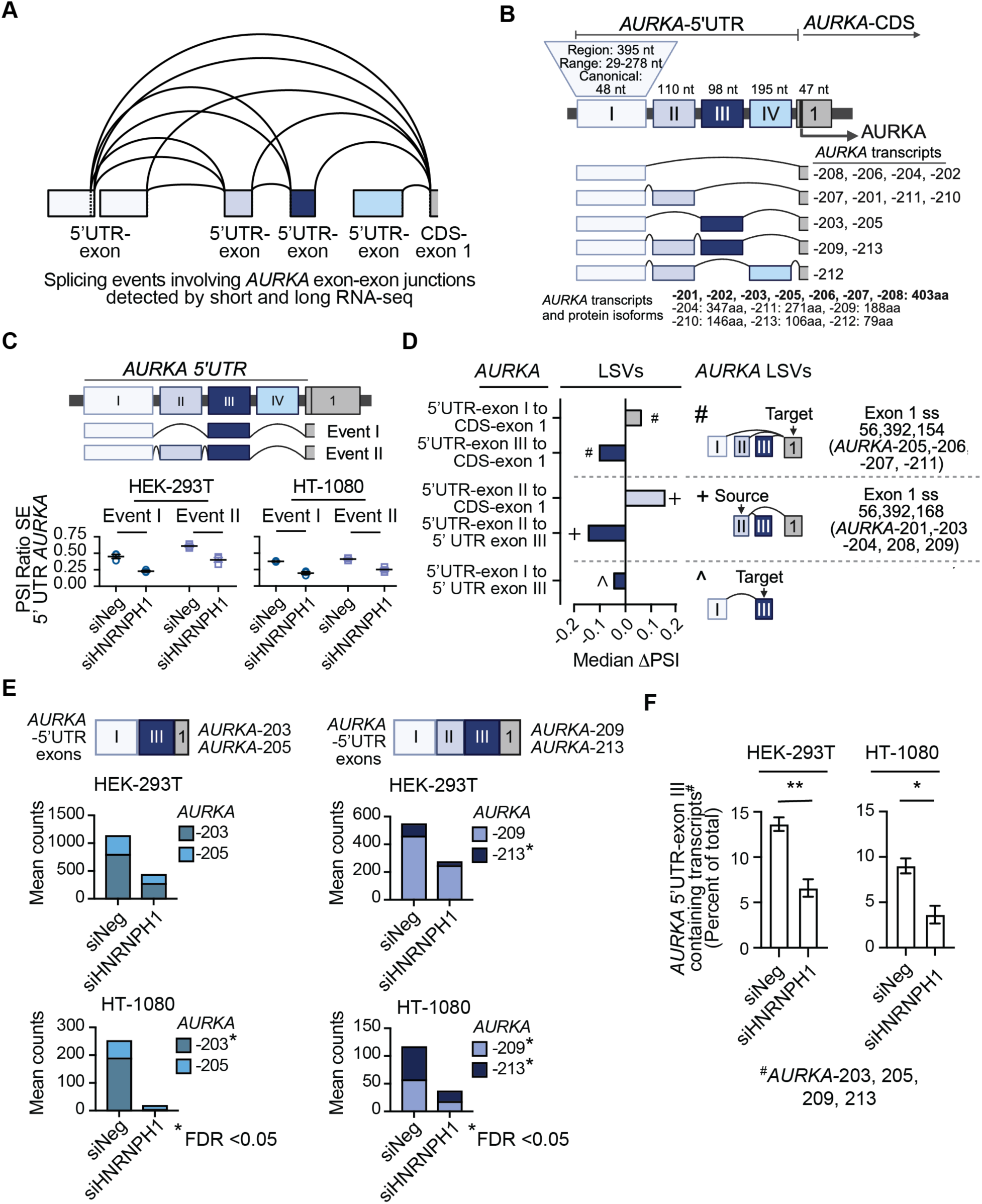
HNRNPH1’s regulation of an *AURKA* 5’UTR alternative splicing event. (**A**) The summary of the exon-exon junctions within the *AURKA* 5’UTR detected using an integrated analysis of short and long-read RNA-seq. Data assessed using the VOILA visualization tool and quantified using the HET quantifier within MAJIQ. (**B**) Schematic model of the *AURKA* 5’ UTR. To distinguish exons corresponding to the regulatory and protein encoding regions of *AURKA* transcripts, we have annotated the *AURKA* 5’ UTR exons using Roman numerals (I-IV). (**C**) The rMATS quantification (PSI ratios) of the events presented in the accompanying schema of *AURKA* transcript variants containing exon III in control (siNeg) and HNRNPH1-depleted (siHNRNPH1) in HEK-293T (left) and HT-1080 (right) cells. (**D**) Analysis of *AURKA-5’UTR*-exon-exon junction usage following the depletion of HNRNPH1 (HET quantifier within MAJIQ). (**E**) Quantification of the indicated *AURKA* transcript variants that exclude *AURKA*-5’UTR-exon III in control (siNeg) and HNRNPH1-depleted (siHNRNPH1) in HEK-293T and HT-1080 cells (EBSeq). (**F**) The percentage of *AURKA* 5’UTR-exon III containing transcript variants in control (siNeg) and HNRNPH1-depleted (siHNRNPH1) cells in the indicated cell lines (students t test, P values * <0.05, ** <0.01).

To quantify the alternative splicing events highlighted by our combined analysis of short- and long-read RNA-seq data, we examined the outputs of the rMATS and MAJIQ analyses. The data from our rMATS analysis identified two events that indicate a decrease in the inclusion of a specific *AURKA-*5’UTR exon III after the silencing of *HNRNPH1* (**Figure 5C**). MAJIQ also quantified a negative ΔPSI for 5’UTR-exon III across three different LSVs with concomitant positive ΔPSI changes for the other exons indicated as affected in the same LSV (**Figure 5D, Supplementary Table S4**). Confirming these findings, differential transcript expression assessed by EBSeq detected decreases in the abundance of individual *AURKA* transcripts that include 5’UTR-exon III (*AURKA*-203, -205, -209, and -213) upon HNRNPH1 depletion, though the significance in the decrease of each respective transcript variant differed between the two cell lines (**Figure 5E**). Moreover, the percent subset abundance analysis of all *AURKA*-5’UTR-exon III containing transcript variants relative to the sum of all protein-coding variants showed an overall significant decrease following HNRNPH1 depletion (**Figure 5F**, **Supplementary Table S4**).

To confirm our RNA-seq analysis, we next designed a PCR based assay to amplify different *AURKA* transcript variants using forward primers that correspond to specific 5’UTR exon I junction sequences and a common reverse primer corresponding to the junction of coding (CDS) exons 1 and 2. In each case, and in both cell lines, we observed reduced inclusion of the *AURKA*-5’UTR exon III following the depletion of HNRNPH1 (**Figures 6A** and **Supplementary S6A**). To complement these findings, we generated a minigene reporter to investigate the effect of *HNRNPH1* silencing on the *AURKA-*5’UTR, taking into consideration variations in the size of *AURKA* 5’UTR-exon I because of the use of alternative splice sites. **Figure 6B** shows the overall organization of the minigene and the transcript-specific PCR assay used to define the use of the indicated exons. The *AURKA-*5’UTR minigene reporter was co-transfected with either a control siRNA (siNeg) or an siRNA targeting *HNRNPH1* (siHNRNPH1) into HEK-293T cells, and 48-hour post-transfection, we used a vector-specific oligonucleotide to synthesize cDNA from the harvested RNA. To detect changes in the usage of *AURKA-*5’UTR exon III, we used a vector-specific reverse and junction-specific forward primers corresponding to different variants of *AURKA*- 5’UTR-exon I (Ia, Ib, and Ic; **Figure 6B**). Using forward primers that overlap the end of each *AURKA*- 5’UTR-exon I variant (Ia, Ib, and Ic) and 5’UTR-exon II, we observed a decrease in the inclusion of *AURKA-*5’UTR-exon III irrespective of the *AURKA*-5’UTR-exon I donor splice site and a concomitant increase in the amplified PCR product that corresponds to the skipping of *AURKA-* 5’UTR-exon III in the samples from the minigene/siHNRNPH1-transfected cells compared to controls (**Figures 6C**). We observed a similar result using forward primers spanning *AURKA*-5’UTR-exon I variant and 5’UTR-exon III junctions (**Figures 6D**). Collectively, these data support that *AURKA-*5’UTR exon III exclusion depends on HNRNPH1 (**Figure 6E**).

**Figure 6:**
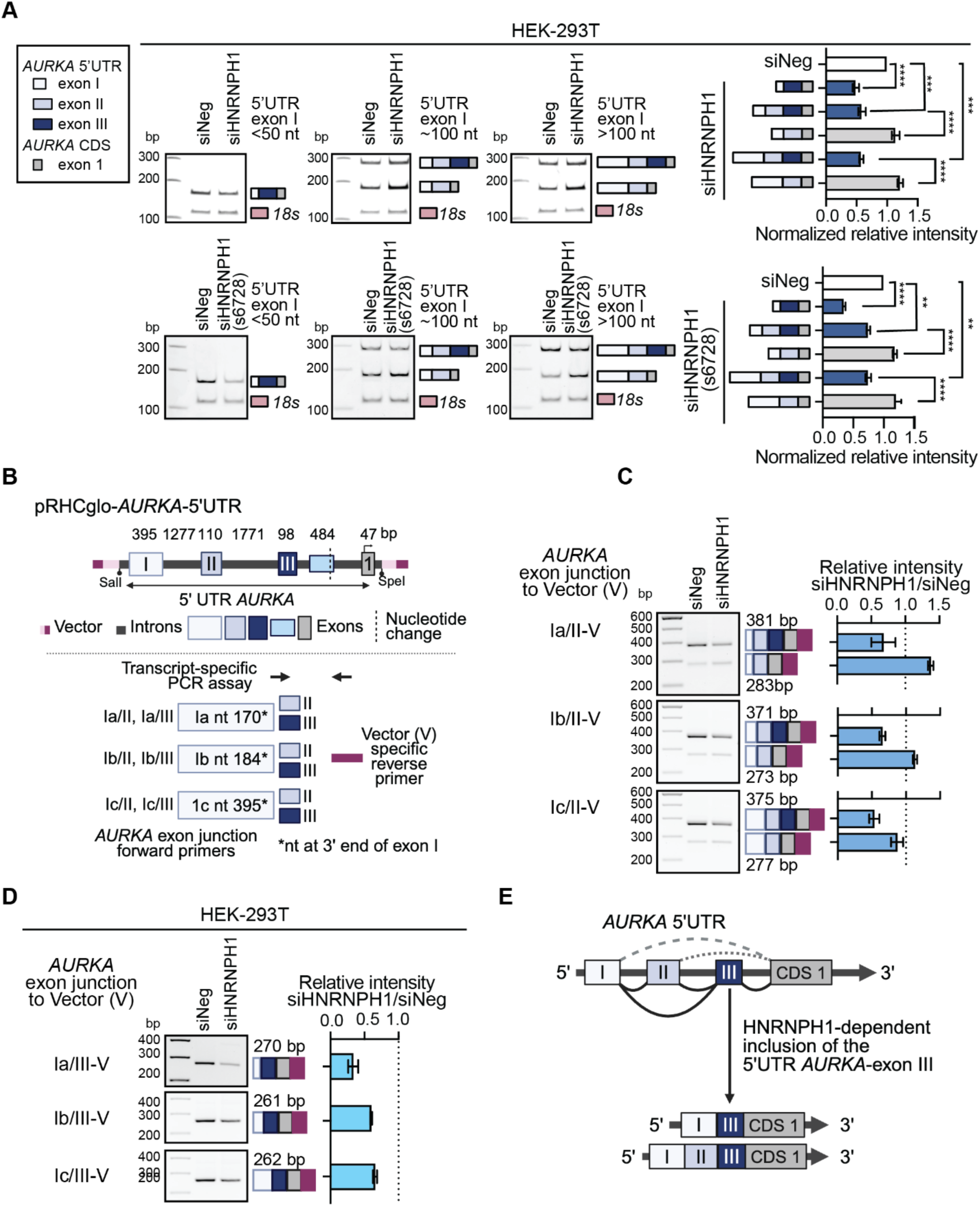
HNRNPH1’s regulation of an *AURKA* 5’UTR skipped exon (SE) event. (**A**) qPCR-based analysis of *AURKA* transcript variants that include *AURKA*-5’UTR-exon III in transfected HEK-293T cells (siNeg and siHNRNPH1: upper panels; siNeg and siHNRNPH1(s6278): lower panels), and quantification of the amplified products normalized to the intensity of the 18S amplified product (mean±SEM of three biological replicates). Statistical analysis: ordinary one-way ANOVA, P values *** <0.001, **** <0.0001. (**B**) Schematic of an *AURKA* minigene encompassing the 5’UTR-exons and CDS exon 1 and adjacent intronic sequences, and the amplification of minigene-derived transcripts. (**C**) Representative images of minigene-derived transcripts amplified using a vector reverse primer and the indicated forward primers specific for the junctions between different lengths of *AURKA* 5’UTR-exon I (a, b, and c) and exon II. The adjacent quantification shows the results of applying this assay to analyze three independent co-transfections. D) Representative images of minigene-derived transcripts amplified using a vector reverse primer and the indicated forward primers specific for the junctions between different lengths of *AURKA* 5’UTR-exon I (a, b, and c) and exon III. The adjacent quantification shows the results of applying this assay to analyze three independent co-transfections. (**E**) A model of HNRNPH1’s proposed regulation of the inclusion of *AURKA* 5’UTR-exon III.

### The inclusion of *AURKA* 5’UTR exon III enhances gene expression

To investigate the effect of including *AURKA* 5’UTR-exon III on gene expression, we next designed a series of luciferase reporters (NanoLuc Luciferase, Promega) to include the *AURKA* minimal promoter (59) and associated sequences that model the different 5’UTRs of at least one *AURKA* transcript variant (**Figure 7A, Supplementary Table S4**). In brief, we designed one set of cassettes to include the small canonical 48 nt version of 5’UTR *AURKA*-exon I (I(S)) and either no additional sequences or 5’UTR *AURKA*-exon sequences upstream of the NanoLuc luciferase reporter gene. The sequence of the I(S) cassette models the 5’UTR of the *AURKA*-208 transcript variant, and I(S)+II, the 5’UTR of *AURKA*-201. The I(S)+III and IS+II+III sequences model the 5’UTR’s of *AURKA*-203 and *AURKA*-209, respectively. The second series of cassettes included a similar arrangement of 5’UTR *AURKA*-exons but utilized a 278 nt longer version of 5’UTR *AURKA*-exon I (I(L)). The I(L) only cassette matches the 5’UTR of *AURKA*-202, I(L)+II, *AURKA*-210, and I(L)+II+III, *AURKA*-213. No reported *AURKA* transcript variant uses I(L) in combination with 5’UTR *AURKA*-exon III (I(L)+III), nor did we detect this theoretical exon organization in our long-read RNA-seq data. To assess the relative strengths of these 5’UTR sequences, we co-transfected the experimental reporters and a plasmid expressing Firefly luciferase into HEK-293T cells and assayed NanoLuc luciferase and Firefly luciferase 48 hours later.

**Figure 7:**
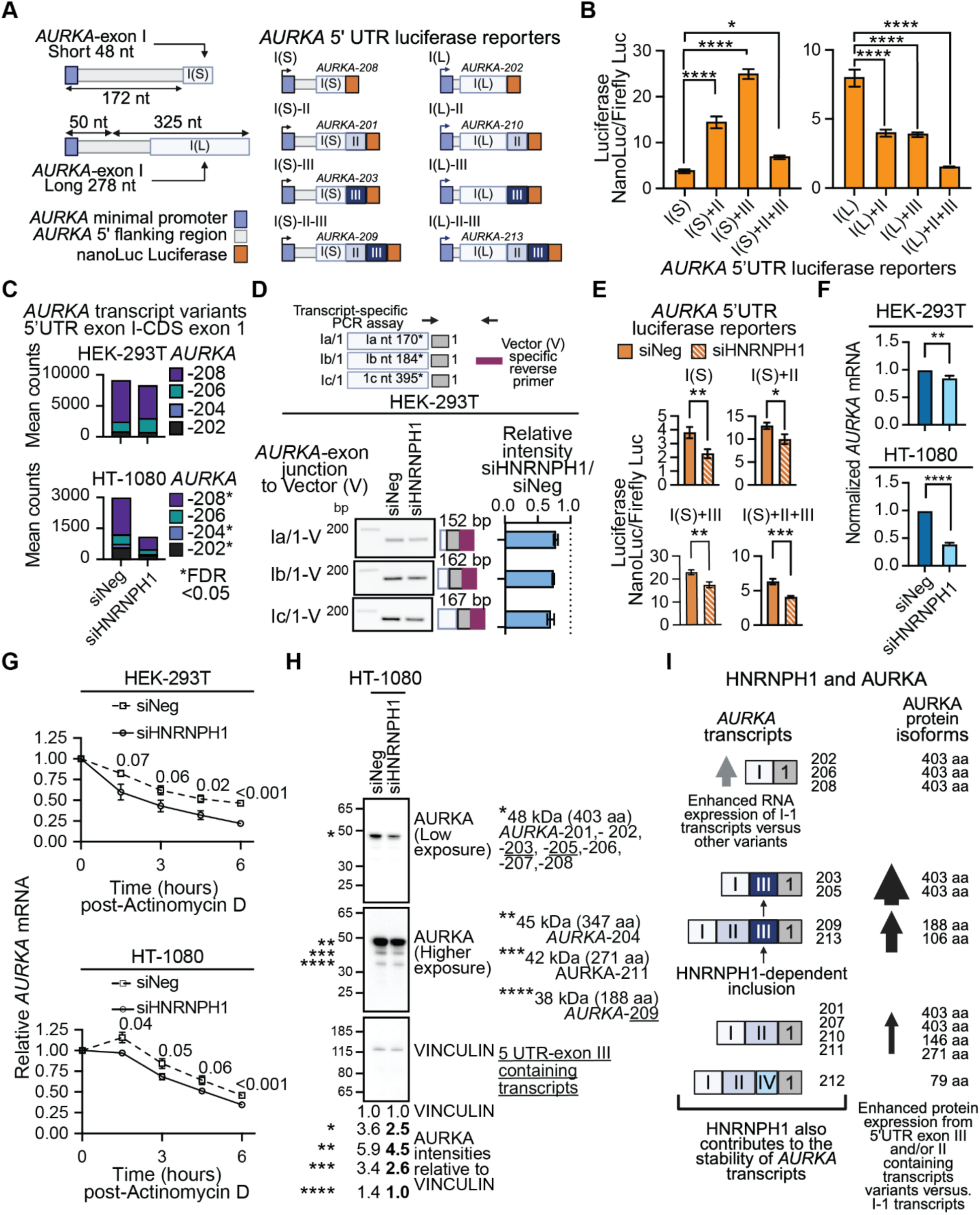
HNRNPH1 regulates the stability of *AURKA* transcripts. (**A**) Schematics of luciferase reporter constructs consisting of a minimal promoter, the indicated *AURKA* 5’UTR sequences, and the NanoLuc luciferase. (**B**) Relative luciferase expression (NanoLuc/Firefly) 48 hours post-transfection of HEK-293T cells with the indicated reporter construct (mean±SEM n=12 per condition). Statistical analysis: ordinary one-way ANOVA, P values *<0.05, ****<0.0001. (**C**) Quantification of the indicated *AURKA* transcript variants expressed in control (siNeg) and HNRNPH1-depleted (siHNRNPH1) in HEK-293T and HT-1080 cells (EBSeq). (**D**) Representative images of a PCR-based amplification of minigene-derived transcripts expressed in control and HEK-293T-HNRNPH1-depleted cells amplified using a vector reverse primer and the indicated forward primers specific for the junctions between different lengths of *AURKA* 5’UTR-exon I (a, b, and c) and CDS exon 1. The adjacent quantification shows the results of applying this assay to analyze three independent co-transfections. (**E**) Relative luciferase expression (NanoLuc/Firefly) 48 hours post-transfection of HEK-293T cells with the indicated reporter construct and the indicated siRNA (mean±SEM n=12 per condition). Statistical analysis: unpaired t test with Welch’s correction, P values * <0.05, ** <0.01, *** <0.001. (**F**) qRT-PCR analysis of *AURKA* expression following the silencing of *HNRNPH1*; unpaired t test with Welch’s correction, P **** <0.001 (see **Supplementary Figure S6C** for quantification of *HNRNPH1* expression). (**G**) qRT-PCR assessment of *AURKA* mRNA at the times indicated post-treatment of siNeg- or siHNRNPH1-transfected HEK-293T and HT-1080 cells with Actinomycin D. P values indicate the statistical difference between the relative levels of *AURKA* at each time point post-addition of Actinomycin D calculated using multiple unpaired t tests. (**H**) The immunoblot analysis of whole cell lysates prepared from siNeg or siHNRNPH1-transfected HT-1080 cells probed using antibodies against the indicated protein. The image shown is representative of three independent transfections per siRNA (see **Supplementary Figure S6D**). The asterisks indicate the detected AURKA protein isoforms, and the values indicate the VINCULIN normalized quantification of each isoform. See **Supplementary Figure S6D** for the analysis of HNRNPH expression. (**I**) A proposed model for the HNRNPH1- dependent alternative splicing of an *AURKA* 5’UTR exon and the stability of AURKA transcripts.

**Figure 7B** shows normalized NanoLuc luciferase levels for each 5’UTR *AURKA* reporter. Interestingly, the longer 278 nt version of *AURKA 5’UTR* exon I generated over twice the mean level of NanoLuc luciferase expression than the canonical 48 nt version. In the context of the long *AURKA 5’UTR* exon I, additional *AURKA 5’UTR* exons decreased luciferase expression, but the addition of *AURKA*-*5’UTR* exon II or exon III to the short *5’UTR* exon I increased luciferase levels significantly, with the mean expression of I(S)+III six times greater than that of the I(S) sequence alone. The dominant *AURKA* transcript variant detected in both HEK-293T and HT-1080 cells corresponds to *AURKA*-208 that utilizes the 48 nt version of *AURKA*-exon I (**Figure 7C**). Of the four *AURKA*-exon I (48 nt) transcript variants that include *AURKA*-exon III, two (AURKA-203 and - 205) encode the same protein as *AURKA*-208, which suggests that the HNRNPH1-dependent inclusion of *AURKA*-exon III could enhance AURKA protein levels via an alternative splicing-dependent mechanism that is independent of transcription.

### HNRNPH1 regulates the stability of *AURKA* transcripts

The complementary minigene and luciferase-based reporter assays we developed to investigate the HNRNPH1-dependent splicing of *AURKA* transcripts revealed one further putative function for HNRNPH1 in regulating *AURKA* gene expression. Specifically, following depletion of HNRNPH1, we observed decreased detection of the *AURKA* 5’UTR-exon I-1 amplified products expressed by our minigene reporter even though expression of these transcripts does not involve an alternative splicing event (**Figure 7D**). Furthermore, following the silencing of *HNRNPH1,* we observed a decrease in the levels of luciferase expressed by all the *AURKA* 5’UTR NanoLuc reporters (**Figure 7E** and **Supplementary Figure S6B**), although expression from the reporter genes is splicing independent. Based on these observations, we hypothesized that non-alternative splicing functions of HNRNPH1 could contribute to the expression of *AURKA* transcripts. To support this hypothesis, we examined the expression of the four *AURKA* transcript variants (AURKA-208, -206, -204, and -202) that utilize only the first 5’UTR *AURKA* exon and exclude the intervening 5’UTR exons (**Figure 7C**). Our short-read RNA-seq data showed that in HT-1080 cells, the silencing of *HNRNPH1* reduces the abundance of three of these four transcript variants, suggesting that HNRNPH1 does contribute to the expression of these transcripts (**Figure 7C, lower panel**). In HEK-293T cells that express higher baseline levels of HNRNPH1, we observed no significant change in the expression of individual *AURKA* transcripts **Figure 7C, upper panel**), but using qRT-PCR, we observed a significant decrease in the overall expression of *AURKA* following depletion of HNRNPH1 in both HEK-293T and HT-1080 cells (**Figure 7F and Supplementary Figure S6C**).

We next assessed if depletion of HNRNPH1 alters the overall stability of *AURKA* transcripts. In brief, we treated HEK-293T and HT-1080 transfected cells (siNeg or siHNRNPH1) with actinomycin-D (ActD) for 1.5, 3, 4.5, and 6.0 hours, harvested RNA at each time point, and assayed *AURKA* levels by qRT-PCR. Both cell lines showed decreased overall stability of *AURKA* in siHNRNPH1 transfected cells versus control, which reached significance at the 6-hour time point (**Figure 7G**). There are limitations to studying the potential impact of these small changes in *AURKA* mRNA variant expression and/or stability at the protein level due to the extremely high expression of AURKA in proliferating cell lines. Nevertheless, in HEK-293T, following depletion of HNRNPH1, we could quantify slight changes in the protein expression in the smaller AURKA isoforms that includes the ∼38 kDa protein encoded by the 5’UTR-exon III containing *AURKA-*209 transcript variant (**Supplementary Figure S6D**). Furthermore, and consistent with the greater response to the depletion of HNRNPH1 observed using HT-1080 cells, we could detect changes in the expression of the dominant 403 amino acid AURKA protein (∼48 KDa) as well as the smaller isoforms in this cell line (**Figure 7H, Supplementary Figure S6D**). Based on these data, we propose a model in which HNRNPH1 regulates the inclusion of an *AURKA* 5’UTR exon that has the potential to enhance protein expression without the need to enhance transcription and contributes to the stability of *AURKA* transcripts overall (**Figure 7I**).

## Discussion

In this study, we present a comprehensive dataset of HNRNPH1-regulated splicing events, initially defined using short-read RNA-seq (**Figure 1**) that we further refined using long-read RNA sequencing. Focusing on specific HNRNPH1-regulated splicing events, we examined the contribution of HNRNPH1 to regulating the expression of proteins that function as: (i) a transcription factor – TCF3 (**Figures 2** and **3**), (ii) a regulator of translation – EIF4G1 (**Figure 4E-Q**), and (iii) a mitotic kinase – AURKA (**Figures 5-7**). In the case of *TCF3* and *EIF4G1,* HNRNPH1 defines the expression of transcript variants encoding different protein isoforms. With respect to the alternative splicing of *AURKA* transcripts, we showed HNRNPH1 regulates the inclusion of a specific 5’UTR exon – 5’UTR *AURKA*-exon III (**Figures 5** and **6**). Critically, luciferase reporter constructs that model the 5’UTR of the most abundant *AURKA* transcript variant, which does not contain this exon, and one that includes the 5’UTR *AURKA*-exon III, showed that inclusion of this exon significantly enhances protein expression (**Figure 7A, B**). We also report that HNRNPH1 depletion alters the splicing of transcripts encoding the adaptor protein NUMB (**Figure 4A, B**) and the RNA binding protein RBM6 (**Figure 4C, D**). Interestingly, the decreased inclusion of *NUMB* exon 12 (CDS exon 9) detected following HNRNPH1 depletion phenocopies the results of silencing *RBM6* on the splicing of *NUMB* (53). Analysis of the *RBM6* long-read RNA seq data showed that HNRNPH1 regulates the inclusion of *RBM6*-exon 6 that encodes a substantial portion of a critical RRM domain leading us to suggest a model in which HNRNPH1 promotes the splicing of *NUMB* transcripts via RBM6 (**Figure 4E**). Collectively, our findings illustrate the importance of HNRNPH1 in regulating the expression of protein-coding genes with critical cellular functions and that by regulating transcript variant expression, HNRNPH1 contributes to the multiplicity of expressed protein isoform expressed, and the tissue-restricted distribution and abundance of specific proteins.

The temporal and spatial expression of bHLH transcription factors and their dimerization with other bHLH proteins, including those categorized as E protein such as TCF3 (E2A), regulate many aspects of development and cell differentiation (reviewed in (60,61). The HNRNPH1-mediated alternative splicing of *TCF3* transcripts (24,25) and this study) that results in the expression of two distinct TCF3 protein isoforms, E12 and E47, adds a further layer of complexity to determining the transcriptional programs regulated by TCF3, which includes those governing the differentiation of hESCs (16,24,25) and B- and T-lymphocyte development (27). In the context of neurogenesis, most studies have focused on the expression of transcripts encoding E47 (i.e., *TCF3*-exon 18b transcripts) and the function of E47 in regulating neuronal cell differentiation (28,62–68). For example, studies using cell-based (neuroblastoma cell lines) (62) and *in vivo* neuronal-specific knockout mouse models (28) have demonstrated that E47 activates the expression of *CDKN1C*, which encodes the p57^Kip2^ cell cycle regulator, and that alterations in E47 function results in impaired neuronal differentiation. In our study, we observed that neuroblastomas exhibit one of the greatest imbalances in the expression of *TCF3* transcripts variants, with *TCF3*-exon 18a transcripts dominating (**Figure 3A**) and that this tumor type also expresses slightly higher levels of *HNRNPH1* than other solid tumor types (**Figure 3B**). These observations could reflect the neural crest origin of NB but prompted by an examination of correlations between MYC activity (defined by a MYC hallmark enrichment score) and *HNRNPH1* and *HNRNPF* expression in different tumor types reported by Chen *et al.,* (19), we evaluated whether the prevalence of MYCN amplification in NB could also be a contributory factor. Though all NB tumors expressed higher levels of *TCF3*- exon 18a transcripts than *TCF3*-exon 18b transcripts, those harboring amplification of MYCN exhibited a greater imbalance (**Figure 3C**), and most MYCN amplified NB cell lines expressed high levels of *TCF3*-18a transcripts (**Figure 3D**). Many of the MYCN positive cell lines also expressed relatively high levels of *HNRNPH1* (**Figure 3E**), leading us to interrogate two independent deposited datasets reporting MYCN’s binding of DNA (48,49) to determine if *HNRNPH1* is a direct target of this oncoprotein. Both datasets showed MYCN binding at the *HNRNPH1* locus (**Figure 3F** and **Supplementary Figure 3D**). These datasets also showed evidence for MYC’s binding of the *HNRNPH1* locus and that other genes encoding members of the HNRNPF/H proteins are targets of MYC and MYCN (**Supplementary Table 3**). Interestingly, the study by Chen *et al*., reported a negative correlation of MYC activity and *HNRNPH1* expression and a positive correlation with HNRNPF in prostate cancer, among others, while they observed the converse correlation in acute myeloid leukemia (AML), that is a positive correlation of MYC activity and *HNRNPH1* expression (19). The complexity of MYC and MYCN’s functions as transcriptional regulators mean it is challenging to connect MYC/MYCN binding and MYC/MYCN activity directly, however, our evaluation of data from NB samples appears more reflective of the positive correlation of MYC activity and HNRNPH1 expression seen in AML. Critically, our findings, together with those of Chen and colleagues, suggest that the proto-oncoproteins MYC/MYCN can impact the expression of HNRNPF/H proteins and thus alternative splicing mechanisms. Our study also suggests that in the context of NB, it will be interesting to assess in future studies whether MYCN’s enhancement of HNRNPH1 expression, and the dominant expression of the E12 encoding *TCF3*- 18a transcript variant influences the function of one of more of the transcription factors that form the CRC associated with NB (44).

Protein synthesis requires the assembly of multiple multi-component translation initiation complexes on RNA. One of these complexes, the EIF4F multi-subunit complex consists of eIf4E, eIF4A, and EIF4G1 (reviewed in (54). Previous studies have described the use of different promoters and alternative start sites to express specific EIF4G1 transcript variants and isoforms (55,56), but the contribution of RNA binding proteins to mediating the expression EIF4G1 is less defined. Here, we show that depletion of HNRNPH1 results in a substantial decrease in the use of the *EIF4G1*-exon 4-5 junction, which is associated with a reduction in the inclusion of canonical 87 nt *EIF4G1*-exon 4 and the adjacent exons, 2 and 3. We also noted decreased expression of transcript variants that include a longer version of *EIF4G1*-exon 4 (147 nt) (*EIF4G1*-209 and -228) or a portion of exon 4 (27 nts) (*EIF4G1*-207). (**Figures 4G-L**). Coupled with these observations is an enhanced detection of transcript variants defined by the skipping of canonical *EIF4G1*-exons 2, 3, and 4, such as *EIF4G1*-206 (**Figures 4L**). These results suggest that mutations affecting *HNRNPH1*, as reported in rare neurodevelopmental disorders (12–14) may not disrupt all EIF4G1 protein expression and thus its function, but it may favor the expression of a non-canonical protein isoform of EIF4G1 that lack residues present in the N-terminus of the full length protein and that are adjacent to the PABP binding domain (**Supplementary Figure S5**). The interaction of polyA- bound PABP and EIF4G1 is critical to facilitating eIF4F’s function as a bridge between the 5’ and 3’ ends of an mRNA. Beyond the initial identification of the PABP binding domain and minimal PABP binding site (58), there is little experimental data assessing the contribution of the amino acids found at the N-terminus of EIF4G1, however, our results suggest that this warrants future study as shifts in the composition of expressed EIF4G1 isoforms may have disease-related implications.

Aurora kinase A (AURKA) is an essential positive regulator of mitosis that exhibits cell-cycle dependent expression, with maximal expression seen in the G2 phase of the cell cycle until mitosis, after which it is present at low levels in G1 that increase during S phase. While the functions of the AURKA protein as a cell-cycle regulator are well defined, the mechanisms that regulate its expression remain less well characterized, particularly the contribution of alternative splicing (69). Here, we confirmed the alternative splicing of *AURKA* pre-mRNAs primarily effects four exons that form the 5’ UTR of *AURKA* transcripts, which here we referred to as 5’ UTR-exons I, II, III and IV (**Figure 5A**, **B**). The first description of the *AURKA* 5’UTR-exons II and III sequences dates from 2000 and involved analysis of *AURKA* (*STK-15*) transcripts expressed in breast cancer cell lines (70). Subsequent studies described mechanisms of translational regulation influenced by the inclusion of specific *AURKA* 5’UTR-exons. In brief, Dobson *et al.* determined that the combination of the longer version of AURKA 5’UTR-exon I and *AURKA* 5’UTR-exon II (which they referred to as the Aurora A 5’ leader) contains an internal ribosome entry site (IRES) element that regulates AURKA protein expression (71). Another study characterized an EGFR responsive element in transcripts within *AURKA* 5’UTR-exon II that can regulate AURKA protein expression, and a subsequent study showed, using a bicistronic reporter assay, that the combination of *AURKA* 5’UTR-exon I (short version) and exon II results in higher reporter protein expression than exon I alone (72,73). Our study confirmed and extended these previous studies to show that the inclusion of *AURKA*-5’UTR exon II or exon III can significantly enhance the levels of protein expression when combined with the canonical *AURKA*-5’UTR exon I of 48 nts and highlighted the dependence of cells on HNRNPH1 to include *AURKA-*5’UTR-exon III. Based on these results, we speculate that the alternative splicing of *AURKA*-5’UTR exons could form part of a transcription-independent mechanism that contributes to the regulation of AURKA protein levels at specific stages of the cell cycle. Future studies will need to evaluate this hypothesis, along with determination of how the combined *AURKA-*5’UTR made up of exons I and III enhances expression compared with *AURKA-* 5’UTR exon I alone. We also observed a reduction in the overall levels and stability of *AURKA* transcripts following the depletion of HNRNPH1. One reason for this observation may relate to HNRNPH1’s interaction with G-rich sequences with the potential to form a G-quadruplex (G4) structure in the presence of a cation (reviewed in (9). Examination of the *AURKA-*5’UTR exon I sequence shows the presence of several G-rich sequences that have the potential to form an RNA G4 (rG4). We and others have highlighted a potential role for HNRNPH1 in inhibiting or destabilizing rG4 formation (22,74), and thus it will be interesting to evaluate in future studies whether HNRNPH1’s interactions with one or more of these G-rich sequences contributes to the stability of *AURKA* mRNAs.

In conclusion, we have focused on selected HNRNPH1-dependent splicing events to (1) interrogate the consequences of MXE or SE events on the expression of specific protein-coding transcript variants and (2) substantiate our transcriptome-wide datasets. Our analysis of specific splicing events highlights how the abundance of HNRNPH1 in cancer or the disruption of its function via mutation could contribute to alterations in the variety and/or abundance of specific protein isoforms. Such findings are critical for furthering our understanding of how changes in alternative splicing mediated by specific RBPs such as HNRNPH1 contribute to genetic and somatic disease.

## Data availability

The data and resources generated are available on reasonable request by contacting the corresponding author. The short and long RNA-seq datasets reported here are available in the Gene Expression Omnibus at GSE298883. See FigShare, DOI: <u>10.6084/m9.figshare.c.7773359</u> for extended data, including gene-specific long-read RNA-seq gff files, complete immunoblot and PCR images, and cell line STR fingerprinting information. See **Supplementary Table S4** for details of the *EIF4G1* GenBank Accessions.

## Author Contributions Statement

Conceptualization: Tayvia Brownmiller, Natasha J. Caplen; Data curation: Tayvia Brownmiller, Erica C. Pehrsson; Formal analysis: Tayvia Brownmiller, Patricio Pichling, Erica C. Pehrsson, Natasha J. Caplen; Investigation: Tayvia Brownmiller, Soumya Sundara Rajan, Tamara Jones, Vernon J. Ebegboni; Methodology: Tayvia Brownmiller, Soumya Sundara Rajan; Resources: Ashish Lal, Ioannis Grammatakakis; Visualization: Tayvia Brownmiller, Natasha J. Caplen; Writing – original draft: Tayvia Brownmiller, Natasha J. Caplen; Writing – review and editing: Tayvia Brownmiller, Soumya Sundara Rajan, Tamara Jones, Erica C. Pehrsson, Natasha J. Caplen; Project administration, supervision, and funding acquisition: Natasha J. Caplen.

## Acknowledgements

We thank the National Cancer Institute (NCI) intramural research program’s CCR Sequencing and CCR Genomic Cores for technical assistance. We thank Javed Khan, Jun S. Wei, and Hsien-Chao Chou (Oncogenomics Section, Genetics Branch, CCR, NCI) for assistance with access to NB cell line and tumor RNA-seq data and NB cell lines. We thank Patrick Zhao and Xinyu Wen for assistance with data deposition. The graphical abstract, schematics of splicing events and reporters, and proposed mechanistic models were created using BioRender.

This research was supported by the Intramural Research Program of the National Institutes of Health (NIH). The contributions of the NIH author(s) were made as part of their official duties as NIH federal employees, are in compliance with agency policy requirements, and are considered Works of the United States Government. However, the findings and conclusions presented in this paper are those of the author(s) and do not necessarily reflect the views of the NIH or the U.S. Department of Health and Human Services.

