## Supplementary Table S1 and Supplementary Figures for "HNRNPH1-mediated splicing events regulate *EIF4G1* transcript variant composition and the organization of the *AURKA* 5’UTR"

**Supplementary Table S1. Reagents and Resources**

**Supplementary Table S2. rMATS analysis, the prioritization of genes for further analysis, TCF3 LSVs and differential transcript expression (EB-seq) (Excel file)**

**Supplementary Table S3. Summary of MYCN and MYC peaks (Excel file)**

**Supplementary Table S4. Gene-specific outputs of long read RNA sequencing (PacBio, Iso-seq), detected LSVs, and differential transcript expression (EB-seq) (Excel file)**

**Supplementary Figure S1. HNRNPH1′s transcriptome-wide regulation of alternative splicing**

**Supplementary Figure S2: HNRNPH1′s regulation of a *TCF3* mutually exclusive splicing event**

**Supplementary Figure S3: The differential expression of *TCF3* transcript variants in neuroblastoma**

**Supplementary Figure S4: HNRNPH1 regulates the inclusion of *NUMB* exon 12 and usage of the *EIF4G1* exon 4-5 junction**

**Supplementary Figure S5: HNRNPH1-regulated splicing events affecting *NUMB* and *EIF4G1* transcripts**

**Supplementary Figure S6: HNRNPH1′s regulation of an *AURKA* 5′UTR skipped exon (SE) event and *AURKA* RNA stability**

### Supplementary Table S1

| Cell lines | Source | STR date |
|---|---|---|
| HEK-293T | ATCC, Manassas, VA | December 2023 |
| HT-1080 | ATCC, Manassas, VA | July 2023 |
| SK-N-FI | Genetics Branch, CCR NCI | November 2024 |
| IMR-32 | Genetics Branch, CCR NCI | November 2024 |

| Primers | Source | Sequence | Comment |
|---|---|---|---|
| *HNRNPH1* forward | Invitrogen (Thermo Fisher Scientific), Waltham, MA | 5′-TACACATGCGGGGATTACCTT-3 | Neckles *et al* (2019) (1) |
| *HNRNPH1* reverse | Invitrogen (Thermo Fisher Scientific) | 5′-CTTCACCAGTTACTCTGCCATC-3' | |
| *NACA* forward | Invitrogen (Thermo Fisher Scientific) | 5′-ACAAGAGCCCTGCTTCAGAT-3′ | |
| *NACA* reverse | Invitrogen (Thermo Fisher Scientific) | 5′-GCTGCTAGTTGTGCTTGCTG-3′ | |
| *RPL27* forward | Invitrogen (Thermo Fisher Scientific) | 5'-GCAAGAAGAAGATCGCCAAG-3' | |
| *RPL27* reverse | Invitrogen (Thermo Fisher Scientific) | 5'-GACGACAGTTTTGTCCAAGG-3' | |
| *TCF3*-exon 17 forward | Invitrogen (Thermo Fisher Scientific) | 5'-AGGAGAAGGAGGACGAGGAG-3' | Yamazaki *et al* (2018) (2) |
| *TCF3*-exon 19 reverse | Invitrogen (Thermo Fisher Scientific) | 5'-GGAGCTGAAAGCACCATCTG-3' | |
| *EIF4G1*-exon 1 forward | Invitrogen (Thermo Fisher Scientific) | 5'-CGCGGACTGAAGGAGACTGA-3' | |
| *EIF4G1*-exon 5 reverse | Invitrogen (Thermo Fisher Scientific) | 5'-AGCAGGGTAGACATGGGCAG-3' | Gonatopoulos-Pournatzis *et al* (2020) (3) |
| *EIF4G1*-exon 5 reverse | Invitrogen (Thermo Fisher Scientific) | 5'-AGCAGGGTAGACATGGGCAG-3' | |
| *AURKA*-exon I/III forward | Invitrogen (Thermo Fisher Scientific) | 5'-TCCTTGGGTCGCAGGCT-3' | Also used for *AURKA* minigene |
| *AURKA*-exon I/II (~100nt) forward | Invitrogen (Thermo Fisher Scientific) | 5'-TCCTTGGGTCGCAGGCT-3' | |
| *AURKA*-exon I/II (>100nt) forward | Invitrogen (Thermo Fisher Scientific) | 5'-CAGCTAGAGGGTCTCACTCC-3' | |
| *AURKA* transcript variant reverse | Invitrogen (Thermo Fisher Scientific) | 5'-GCCTTAACAGGTCCTGAAATGC-3' | |
| 18s forward | Invitrogen (Thermo Fisher Scientific) | 5'-GTAACCCGTTGAACCCCATT-3' | |
| 18s reverse | Invitrogen (Thermo Fisher Scientific) | 5'-GGACTTAATCAACGCAAGCT-3' | |
| RTRHC Reverse (cDNA synthesis) | Invitrogen (Thermo Fisher Scientific) | 5'-GAGCTTTGCAGCAACAGTAAC-3' | Singh and Cooper (2006) (4) |
| *AURKA*-exon Ia forward | Invitrogen (Thermo Fisher Scientific) | 5'-TTGGGTCGCAGGCATCATG-3' | |
| *AURKA*-exon Ia-II forward | Invitrogen (Thermo Fisher Scientific) | same as *AURKA*-exon I/II (~100nt) | |
| *AURKA*-exon Ia-III forward | Invitrogen (Thermo Fisher Scientific) | same as *AURKA*-exon I/III | |
| *AURKA*-exon Ib forward | Invitrogen (Thermo Fisher Scientific) | 5'-ACGGGCATCATGGACCGAT-3' | |
| *AURKA*-exon Ib-II forward | Invitrogen (Thermo Fisher Scientific) | 5'-GACGGGGTCTCACTCCATT-3' | |

| | | | |
|---|---|---|---|
| *AURKA*-exon Ib-III forward | Invitrogen (Thermo Fisher Scientific) | 5'-GACGGGCTGGAGTGCAAT-3' | |
| *AURKA*-exon Ic forward | Invitrogen (Thermo Fisher Scientific) | 5'-CAGCTAGAGGCATCATGGACC-3' | |
| *AURKA*-exon Ic-II forward | Invitrogen (Thermo Fisher Scientific) | same as AURKA-exon I/II (>100nt) | |
| *AURKA*-exon Ic-III forward | Invitrogen (Thermo Fisher Scientific) | 5'-CTAGAGGCTGGAGTGCAATGG-3' | |
| RTRHC Reverse (minigene) | Invitrogen (Thermo Fisher Scientific) | 5'-GGAATATGCACCACTTGGAACA-3' | |
| *AURKA* (qPCR) forward | Invitrogen (Thermo Fisher Scientific) | 5'-GGAATATGCACCACTTGGAACA-3' | CDS-exon 5 |
| *AURKA* (qPCR) reverse | Invitrogen (Thermo Fisher Scientific) | 5'-TAAGACAGGGCATTTGCCAAT-3' | CDS-exon 6 |
| *GAPDH* forward | Invitrogen (Thermo Fisher Scientific) | 5'-TGCACCACCAACTGCTTAGC-3' | Actinomycin-D qPCR |
| *GAPDH* reverse | Invitrogen (Thermo Fisher Scientific) | 5'-GGCATGGACTGTGGTCATGAG-3' | |

| **PCR/qPCR Reagents** | **Company** | **Catalog #** |
|---|---|---|
| 5X iScript RT Supermix (cDNA synthesis) | Bio-Rad Laboratories, Hercules, CA | 1708841 |
| SuperScript II Reverse Transcriptase (cDNA synthesis) | Invitrogen (Thermo Fisher Scientific) | 18064-014 |
| Q5 Hot Start High-Fidelity 2X Master mix | New England BioLabs, Ipswich, MA | M0494S |
| Phusion High-Fidelity PCR Master mix with GC Buffer | New England BioLabs | M0532S |
| PowerUp SYBR Green Master Mix | Applied Biosystems (Thermo Fisher Scientific) | A25742 |
| TrackIt 100bp DNA ladder | Invitrogen (Thermo Fisher Scientific) | 10488058 |
| 6% TBE Gel | Invitrogen (Thermo Fisher Scientific) | EC6265BOX |
| Novex Hi-Density TBE Sample Buffer (5X) | Invitrogen (Thermo Fisher Scientific) | LC6678 |
| 100bp DNA Ladder | Invitrogen (Thermo Fisher Scientific) | 15628019 |
| SYBR Safe DNA Gel Stain | Invitrogen (Thermo Fisher Scientific) | S33102 |

| **Reporter plasmids and DNA cassettes** | **Source** | **Catalog number or sequence** |
|---|---|---|
| pRHCglo | Addgene, Watertown, MA | 80169; Singh and Cooper (2006) |
| *AURKA*-5'UTR DNA cassette | Custom synthesis Genewiz, South Painfield, NJ<br><br>Restriction enzymes: **SalI (gtcgac)** and **SpeI (actagt)** | **gtcgac**ggttcttagggagcaagtgcgcctgcgcggtgtgcgcccttaaacgcgactcaaggcgt<br>cgggtttgttgtcaaccaatcacaaggcagcctcgctcgagcgcaggccaatcggctttctagctagag<br>ggtttaactcctatttaaaaagaagaacctttgaattctaacggctgagctcttggaagacttgggtcctt<br>gggtcgcaggtgggagccgacgggtgggtagaccgtgggggatatctcagtggcggacgaggac<br>ggcggggacaaggggcggctggtcggagtggcggagcgtcaagtcccctgtcggttcctccgtccc<br>tgagtgtccttggcgctgccttgtgcccgcccagcgcctttgcatccgctcctgggcaccgaggcgccc<br>tgtaggatactgcttgttacttattacagctagaggtacaagggggtttgttgagtggtgttgacatgcgc<br>gggaggggtgggtgggcttcagattggattttgtcctccgagatcacctgggtaagaaaagcacaga<br>aggtctctgctgtgttggagttcacgttctagtgggtggggagacagacggtaagcgtacggagaac<br>ttgcagctggggtgggtgttacagaggaaaagcaggagtgcggtttaacgggggccgctttagata<br>gaatagcctaagaaggcccttgtcctggctggatgagtgggtgaattgatgaatgagaacctccttgc<br>agaggccttcccggtccgtacttctcagaccgtagaccttggagcagttgtaccgggataacttgcagt<br>gcttgttaaacatatagctccaccgagagcccggtaaatcagaatctctgaggggtaggacccaggtg<br>tctagttattaacaaactctccagatgattcttaggtttcattgagaactagggatgcagaccggtgcata<br>caaatcgtctggggacgttaaaatgcagaatctgtttttttagacctggggttaggcctcagagtctccat<br>ttctaagaaccatccggctgaggtagacgtagccactgtccttaactctcgtaatgtctcttcctcttccgt<br>aaccttccttgtcccttgaattaaacgtttttcagcaacctactcagttcgtccttcccttcatctctgcagaca |

| | | |
|---|---|---|
| | | tgcacaggtctgagggaggaaggaataaaccgtataaacctcctgcgctattagcctaacagcttttct<br>attcaaaatagtaggacttctggtttgaactgaatggatcctgtgaaagtcatctggttcttcaatctggtt<br>gcagattagagtcacctgaagggcagtctccttggcgttgtctccagaattctggattagaatcttattcc<br>attctgcttgttattcaatttccctagaaagaaaggtagaataaattggagcaaatgcctgtagcttctgtc<br>agaagaatgttgaataaatgttgttaggcctatgtgatctcattagactgctacttagaattgtaaggga<br>agtaaagcattagagcatgtgtgaaattaaatatttgattaacacaagtgtgcatttccttgttgctgtttat<br>caacttttacttacccactgtttttttataagggctgcagcctgtagtctgggcctggcttcatcatggaatt<br>atttgcttaattgtaaaatggtaatcttaatttttttttttgagacagggtctcactccattgcccaggccaga<br>gtgcggggatatttgataagaaacttcagtgaaggccgggcgcggtggctcatgcccgtaatcccag<br>cattttcggaggccgaggtgggtggatcacctgaggtcgggagttccagaccagccctaccaacata<br>gagaaaccctgtctctactaaaaatacaaaatcagctgggcgtggtggtgcatgtctgtaatcccagct<br>actctggaggctgaggcaggagaattgcttgaacccaggaggtggaggttgcgggtgagccgaga<br>ttgcgccattgcactccagcctgggcaacaagagcaaactctatctcaaaaaaaaaaaaaaagaaacct<br>cagtgaaatgaaagactgaactacacacctgggggaaaagcgtaataaagggagcaagaagtgca<br>aatgcttatggcaggggacttatttgcagaaagaaagccttgtgtggctggtacacggttgacaagaa<br>gaggagatgaagtcagagaggtagttagggtccatgttctgtgagacttcaataggcctgtctatgca<br>ggttaggagtttgcattgttttgattgagttgggagccatggggtcttctgaactgaggatcaacacat<br>ctgatttgttaatgagattgccggcctcagtgttaagaatggaccatgacagggtaagggtggaagtg<br>gggaggccattgggaggctgttggaaaaaactcaagggagagagaataatggtttgaccaggtggt<br>agtagtagaggtggtgagcagtgatcttgaaggcagagcctacagaatatgctggtggattaaatga<br>agggtttgcgagagagaacctcgtaaagggttattatgccaaggcttttggcctgagcaactgacagg<br>atgggtttgccatgcactgagatgaggatgtctagagactagatttggggaggtgagagtggggag<br>agattaggacttctgtttgggacatgttagaatggagatgccgctcagataatcaggtgcagatgttgc<br>atagaaagtagaaaatgtagatcaagagttcaggggagaggtctgggctgaagaatgaatatttggg<br>agtcctcagcctgtagatggtactgaaagcaaagagactggatgtactcggctagggagtaattgga<br>gatggagaagtcattctaggcctgagctcagggcacacccatgaacagggaggtgagaagtggag<br>aagggaagaggttcagttttggatatgttaggattgaggtgcctggcagacatttttgggagatcctgg<br>gcaaatccttggatatccaagtctgacagtccaaggagagactagggcagaagacgcactttgagtg<br>gtcctcatgaaggcatttagagctgaggcactgggtgatatggatccctggtgaaggtgtagatgga<br>gaagtggagggtccagggactcgttcttgagtcaccccagacccttgatctggcaaaatgagactga<br>cccagcaaggagggaaagacaggacgactaggtggtagataaacctggaagtgaagtgaagaaa<br>aaagtgtaagatgtgatcaaccttgtcagattctgctgaggatgcaagtacggggaaggtggaattga<br>cccttgggtttggcagcagtaggtgtcagccatgagcctgacaatgtggtggagacaaaaaagccag<br>gtggggatgggctcaggggaggggagaggagcagtagaggtggcaagtgtggacggttcttctg<br>gagctttgctgttggagaggagggggtcattgttagcaacgataatagaataaagaggacttttatttt<br>attttattgttttgagacgaagtctcgctcttttcccataggctggagtgcaatggtgtgatctcagctcact<br>gcaacctctgcttcctgggtttaagtgattctcctgcctcagcctcccgagtagctgggattacaggtgac<br>tgccaccacgctcagctaagttttgtatttttagtagagacggggtttcaccatgttggccaggctggtct<br>cgaactcttgacctcaggtgatccgcccacctcggcctcccaaaatgctgggattacaggtgtaagcca<br>ccatgcccggcctaaagaggactttttaaggtagcaggtatgccagtaggctgcacagggatggaa<br>gtgatccattagaggagagatgatggagcagcaagaccatcaggagtgaagactccgtgacagagg<br>atgggacgcagtgcccaaggggaggttggccttgcagggaaagcacatgcctgtccctccctcataa<br>gcttcatttggacaaaacatgtaaaatccggtgtgttgtggaggccttttgattggggaactgtaacgct<br>gcctatcgagcaacagcactttaagcaggtggctttgttcaaattaaaggttcttctttttcttttcaggcat<br>catggaccgatctaaagaaaactgcatttcaggacctgttaaggtaaattgaataatctgtaatctcattc<br>acatttataaac**actagt** |
| pGL4.54[luc2/TK] | Promega, Madison, WI | E5061 |
| pNL2.2[NlucP/Hygro] | Promega | N1071 |
| *AURKA*-exon I (S) | Genewiz **NheI** (**gctagc**) and **HindIII (aagctt)** | **gctagc**accacttccgggttcttagggagcaagtgcgcctgcgcgcggtgtgcgcccttaaacgcg<br>actcaaggcgtcgggtttgttgtcaaccaatcacaaggcagcctcgctcgagcgcaggccaatcggc<br>tttctagctagagggtttaactcctatttaaaaagaagaacctttgaattctaacggctgagctcttgga<br>agacttgggtccttgggtcgcag**aagctt** |
| *AURKA*-exon I (S)+II | Genewiz | **gctagc**accacttccgggttcttagggagcaagtgcgcctgcgcgcggtgtgcgcccttaaacgcg<br>actcaaggcgtcgggtttgttgtcaaccaatcacaaggcagcctcgctcgagcgcaggccaatcggc<br>tttctagctagagggtttaactcctatttaaaaagaagaacctttgaattctaacggctgagctcttgga<br>agacttgggtccttgggtcgcaggctggagtgcaatggtgtgatctcagctcactgcaacctctgcttc<br>ctgggtttaagtgattctcctgcctcagcctcccgagtagctgggattacag**aagctt** |

| | | |
|---|---|---|
| *AURKA*-exon I (S)+III | Genewiz | **gctagc**accacttccgggttcttagggagcaagtgcgcctgcgcgcggtgtgcgcccttaaacgcg<br>actcaaggcgtcgggtttgttgtcaaccaatcacaaggcagcctcgctcgagcgcaggccaatcggc<br>tttctagctagagggtttaactcctatttaaaaagaagaacctttgaattctaacggctgagctcttgga<br>agacttgggtccttgggtcgcaggctggagtgcaatggtgtgatctcagctcactgcaacctctgcttc<br>ctgggtttaagtgattctcctgcctcagcctcccgagtagctgggattacag**aagctt** |
| *AURKA*-exon I (S)+II+III | Genewiz | **gctagc**accacttccgggttcttagggagcaagtgcgcctgcgcgcggtgtgcgcccttaaacgcg<br>actcaaggcgtcgggtttgttgtcaaccaatcacaaggcagcctcgctcgagcgcaggccaatcggc<br>tttctagctagagggtttaactcctatttaaaaagaagaacctttgaattctaacggctgagctcttgga<br>agacttgggtccttgggtcgcagggtctcactccattgcccaggccagagtgcggggatatttgataa<br>gaaacttcagtgaaggccgggcgcggtggctcatgcccgtaatcccagcattttcggaggccgagg<br>ctggagtgcaatggtgtgatctcagctcactgcaacctctgcttcctgggtttaagtgattctcctgcctc<br>agcctcccgagtagctgggattacag**aagctt** |
| *AURKA*-exon I (L) | Genewiz | **gctagc**accacttccgggttcttagggagcaagtgcgcctgcgcgcggtgtgcgcccttaaacgcg<br>actcaaggcgtcgggtttgttgtcaaccaatcacaaggcagcctcgctcgagcgcaggccaatcggc<br>tttctagctagagggtttaactcctatttaaaaagaagaacctttgaattctaacggctgagctcttgga<br>agacttgggtccttgggtcgcaggtgggagccgacgggtgggtagaccgtgggggatatctcagt<br>ggcggacgaggacggcggggacaaggggcggctggtcggagtggcggagcgtcaagtcccct<br>gtcggttcctccgtccctgagtgtccttggcgctgccttgtgcccgcccagcgcctttgcatccgctcct<br>gggcaccgaggcgccctgtaggatactgcttgttacttattacagctagag**aagctt** |
| *AURKA*-exon I (L)+II | Genewiz | **gctagc**accacttccgggttcttagggagcaagtgcgcctgcgcgcggtgtgcgcccttaaacgcg<br>actcaaggcgtcgggtttgttgtcaaccaatcacaaggcagcctcgctcgagcgcaggccaatcggc<br>tttctagctagagggtttaactcctatttaaaaagaagaacctttgaattctaacggctgagctcttgga<br>agacttgggtccttgggtcgcaggtgggagccgacgggtgggtagaccgtgggggatatctcagt<br>ggcggacgaggacggcggggacaaggggcggctggtcggagtggcggagcgtcaagtcccct<br>gtcggttcctccgtccctgagtgtccttggcgctgccttgtgcccgcccagcgcctttgcatccgctcct<br>gggcaccgaggcgccctgtaggatactgcttgttacttattacagctagagggtctcactccattgccc<br>aggccagagtgcggggatatttgataagaaacttcagtgaaggccgggcgcggtggctcatgccc<br>gtaatcccagcattttcggaggccgag**aagctt** |
| *AURKA*-exon I (L)+III | Genewiz | **gctagc**accacttccgggttcttagggagcaagtgcgcctgcgcgcggtgtgcgcccttaaacgcg<br>actcaaggcgtcgggtttgttgtcaaccaatcacaaggcagcctcgctcgagcgcaggccaatcggc<br>tttctagctagagggtttaactcctatttaaaaagaagaacctttgaattctaacggctgagctcttgga<br>agacttgggtccttgggtcgcaggtgggagccgacgggtgggtagaccgtgggggatatctcagt<br>ggcggacgaggacggcggggacaaggggcggctggtcggagtggcggagcgtcaagtcccct<br>gtcggttcctccgtccctgagtgtccttggcgctgccttgtgcccgcccagcgcctttgcatccgctcct<br>gggcaccgaggcgccctgtaggatactgcttgttacttattacagctagaggctggagtgcaatggt<br>gtgatctcagctcactgcaacctctgcttcctgggtttaagtgattctcctgcctcagcctcccgagtagc<br>tgggattacag**aagctt** |
| *AURKA*-exon I (L)+II+III | Genewiz | **gctagc**accacttccgggttcttagggagcaagtgcgcctgcgcgcggtgtgcgcccttaaacgcg<br>actcaaggcgtcgggtttgttgtcaaccaatcacaaggcagcctcgctcgagcgcaggccaatcggc<br>tttctagctagagggtttaactcctatttaaaaagaagaacctttgaattctaacggctgagctcttgga<br>agacttgggtccttgggtcgcaggtgggagccgacgggtgggtagaccgtgggggatatctcagt<br>ggcggacgaggacggcggggacaaggggcggctggtcggagtggcggagcgtcaagtcccct<br>gtcggttcctccgtccctgagtgtccttggcgctgccttgtgcccgcccagcgcctttgcatccgctcct<br>gggcaccgaggcgccctgtaggatactgcttgttacttattacagctagagggtctcactccattgccc<br>aggccagagtgcggggatatttgataagaaacttcagtgaaggccgggcgcggtggctcatgccc<br>gtaatcccagcattttcggaggccgaggctggagtgcaatggtgtgatctcagctcactgcaacctct<br>gcttcctgggtttaagtgattctcctgcctcagcctcccgagtagctgggattacag**aagctt** |

| **Restriction Enzymes & Cloning** | **Company** | **Catalog #** |
|---|---|---|
| PstI | New England BioLabs | R3140S |
| SalI | New England BioLabs | R3138S |
| SpeI | New England BioLabs | R3133S |
| FastDigest NheI | Thermo Fisher Scientific | FD0974 |
| FastDigest HindIII | Thermo Fisher Scientific | FD0505 |
| T4 DNA Ligase | New England BioLabs | M0202S |
| T4 Polynucleotide Kinase | New England BioLabs | M0201S |
| Quick CIP | New England BioLabs | M0525S |
| One Shot MAX Efficiency DH5α-T1R Competent Cells | ThermoFisher Scientific | 12297016 |

| Antibodies/Stain | Source | Catalog # | Assay Concentration |
|---|---|---|---|
| Rabbit anti-AURKA | Cell Signaling Technologies, Danvers, MA | 14475 | Immunoblot 1:500 |
| Rabbit anti-HNRNPH | Bethyl Laboratories, Montgomery, TX | A300-511A | Immunoblot 1:1500 |
| Mouse anti-VINCULIN | Santa Cruz Biotechnology, Dallas, TX | sc-25336 | Immunoblot 1:1500 |
| Rabbit anti-ACTIN | Cell Signaling Technologies, | 4967 | Immunoblot 1:1500 |
| Rabbit anti-GAPDH | Abcam | ab8245 | Immunoblot 1:1500 |
| Rabbit anti-N-MYC | Cell Signaling Technologies | 9405S | Immunoblot 1:1000 |
| Goat Anti Rabbit IgG-HRP | Cell Signaling Technologies | 7074S | Immunoblot 1:3000 |
| Goat Anti Mouse IgG-HRP | Cell Signaling Technologies | 7076S | Immunoblot 1:3000 |

| siRNAs | Company | Catalog # | Sequence |
|---|---|---|---|
| siNeg | Qiagen, Germantown, MD | SI03650318 | - |
| siHNRNPH1 (s6730) | Ambion (Thermo Fisher Scientific) | s6730 | 5'-GGAUUUGGGUCAGAUAGAUTT-3' |
| siHNRNPH1 (s6728) | Ambion (Thermo Fisher Scientific) | s6728 | 5'-GGAAGCAUACUGGUCCAAAUTT-3' |

| Chemicals and other resources | Company | Catalog # |
|---|---|---|
| Fetal Bovine Serum | Invitrogen (Thermo Fisher Scientific) | A56708-01 |
| DMEM | Invitrogen (Thermo Fisher Scientific) | 11995-065 |
| EMEM | ATCC | 30-2003 |
| RPMI-1640 | Invitrogen (Thermo Fisher Scientific) | 11875-093 |
| DPBS | Invitrogen (Thermo Fisher Scientific) | 10010-023 |
| 0.25% Trypsin-EDTA | Invitrogen (Thermo Fisher Scientific) | 25200-056 |
| MycoAlert Mycoplasma Detection Kit | Lonza, Walkersville, MD | LT07-218 |
| Plasmocin | InvivoGen USA, San Diego, CA | ant-mpp |
| Lipofectamine RNAiMAX | Invitrogen (Thermo Fisher Scientific) | 13778100 |
| Lipofectamine 2000 | Invitrogen (Thermo Fisher Scientific) | 11668500 |
| OptiMEM | Invitrogen (Thermo Fisher Scientific) | 31985-070 |
| Maxwell 16 LEV simplyRNA purification kit | Promega, Madison, WI | AS1390 |
| Nano-Glo Dual-Luciferase Reporter Assay System | Promega | N1610 |
| Actinomycin D | Sigma, St. Louis, MO | A1410 |
| DMSO | Invitrogen (Thermo Fisher Scientific) | D12345 |
| QIAquick Gel Extraction Kit | Qiagen | 28704 |
| QIAquick PCR Purification Kit | Qiagen | 28104 |
| PureLink HiPure Plasmid Midiprep Kit | Invitrogen (Thermo Fisher Scientific) | K210005 |
| PureLink HQ Mini Plasmid Purfication Kit | Invitrogen (Thermo Fisher Scientific) | 2385512 |
| Cell Extraction Buffer | Invitrogen (Thermo Fisher Scientific) | FNN0011 |
| Pierce BCA Protein Assay Kit | Thermo Fisher Scientific | A55864 |
| Poly-L-Lysine | Millipore Sigma, Burlington, MA | P8920 |
| PureCol | Advanced Biomatrix, Carlsbad, CA | 5005 |

| | | |
|---|---|---|
| Triton-X 100 | Millipore Sigma | X100 |
| NuPAGE™ Bis-Tris Mini Protein Gels, 4–12% | Thermo Fisher Scientific | NP0322BOX |
| NuPAGE™ MOPS SDS Running Buffer (20X) | Thermo Fisher Scientific | NP0001 |
| PageRuler™ Prestained Protein Ladder, 10 to 180 kDa | Thermo Fisher Scientific | 26616 |
| Cell Line Nucleofector® Kit V | Lonza | VCA-1003 |
| Cell Line Nucleofector® Kit L | Lonza | VCA-1005 |
| Cell Line Nucleofector® Kit R | Lonza | VCA-1001 |

| **Software, R packages, and other tools** | |
|---|---|
| GraphPad Prism version 10.2.3 | https://www.graphpad.com/ |
| Microsoft Excel (Mac Version 16.95.4) | https://www.microsoft.com/en-us/microsoft-365/excel |
| CCBR pipeliner Renee | https://ccbr.github.io/RENEE/latest/#renee---rna-sequencing-analysis-pipeline |
| R/R studio | https://www.r-project.org/ https://posit.co/download/rstudio-desktop/ |
| ImageJ | https://imagej.net/software/fiji/ |
| rMATS | https://github.com/Xinglab/rmats-turbo |
| Maser | https://bioconductor.org/packages/release/bioc/html/maser.html |
| Metascape | https://metascape.org/gp/index.html#/main/step1 |
| MAJIQ and MAJIQ-L | https://majiq.biociphers.org/ |
| MAJIQlopedia | https://majiq.biociphers.org/majiqlopedia/ |
| EBSeq | https://bioconductor.org/packages/release/bioc/html/EBSeq.html |
| IsoSeq4 | https://isoseq.how/ |
| Target/TCGA data: UCSC Xena | https://xenabrowser.net/ |

| **PCR Conditions** | | | |
|---|---|---|---|
| ***TCF3*-exon 18 Splice Assay PCR (2 µL cDNA; Phusion GC)** | | | |
| 98°C for 30 sec | | | |
| 98°C for 10 sec | 35 cycles | | |
| 64°C for 10 sec | | | |
| 72°C for 30 sec | | | |
| 72°C for 5 min | | | |
| ***EIF4G1* (exon 1 to exon5 PCR; 2 µL cDNA; Phusion GC)** | | | |
| 98°C for 30 sec | | | |
| 98°C for 8 sec | 35 cycles | | |
| 66°C for 12 sec | | | |
| 72°C for 15 sec | | | |
| 72°C for 5 min | | | |
| ***EIF4G1* (exon 4/µexon PCR; 2 µL cDNA: Phusion GC)** | | | |
| 98°C for 30 sec | | | |
| 98°C for 8 sec | 35 cycles | | |
| 65°C for 12 sec | | | |
| 72°C for 15 sec | | | |
| 72°C for 5 min | | | |
| ***AURKA* and 18s (endogenous transcript variants; 1 µL cDNA; Q5 Hot Start)** | | | |
| 98°C for 30 sec | | | |
| 98°C for 8 sec | 30 cycles | | |
| 65°C for 20 sec | | | |
| 72°C for 8 sec | | | |
| 72°C for 2 min | | | |
| **AURKA Minigene** | | | |
| Junction specific forward primer | cDNA (µL) | Annealing temperature (°C) | # of Cycles |
| Ia+II | 8 | 65 | 35 |
| Ia+III | 8 | 66 | 35 |

| | | | |
|---|---|---|---|
| Ia+CDS1 | 8 | 66 | 35 |
| Ib+II | 4 | 64 | 32 |
| Ib+III | 4 | 65 | 32 |
| Ib+CDS1 | 4 | 66 | 32 |
| Ic+II | 4 | 64 | 32 |
| Ic+III | 4 | 65 | 32 |
| Ic+CDS1 | 4 | 65 | 32 |

| Tumor data extracted from MAJIQlopedia (5) | | |
|---|---|---|
| Abbreviation | Tumor type | Sample number and original data source |
| ACC | Adrenocortical carcinoma | n=153, TCGA |
| BLCA | Bladder urothelial carcinoma | n=432, TCGA |
| BRCA | Breast invasive carcinoma | n=498, TCGA |
| CCSK | Clear Cell Sarcoma of the Kidney | n=13, TARGET |
| CESC | Cervical squamous cell carcinoma and endocervical adenocarcinoma | n=309, TCGA |
| CHOL | Cholangiocarcinoma | n=35, TCGA |
| COAD | Colon adenocarcinoma | n=352, TCGA |
| ESCA | Esophageal carcinoma | n=181, TCGA |
| GBM | Glioblastoma multiforme | n=158, TCGA |
| HNSC | Head and neck squamous cell carcinoma | n=565, TCGA |
| KICH | Kidney chromophobe | n=155, TCGA |
| KIRC | Kidney renal clear cell carcinoma | n=298, TCGA |
| KIRP | Kidney renal papillary cell carcinoma | n=323, TCGA |
| LGG | Brain lower grade glioma | n=499, TCGA |
| LIHC | Liver hepatocellular carcinoma | n=423, TCGA |
| LUAD | Lung adenocarcinoma | n=500, TCGA |
| LUSC | Lung squamous cell carcinoma | n=496, TCGA |
| MESO | Mesothelioma | n=166, TCGA |
| NB | Neuroblastoma | n=168, TARGET |
| OV | Ovarian Serous cystadenocarcinoma | n=401, TCGA |
| PCPG | Pheochromocytoma and Paraganglioma | n=187, TCGA |
| PAAD | Pancreatic adenocarcinoma | n=183, TCGA |
| PRAD | Prostate adenocarcinoma | n=500, TCGA |
| READ | Rectum adenocarcinoma | n=104, TCGA |
| SARC | Sarcoma | n=263, TCGA |
| SKCM | Skin cutaneous melanoma | n=473, TCGA |
| STAD | Stomach adenocarcinoma | n=374, TCGA |
| TGCT | Testicular germ cell tumors | n=139, TCGA |
| THCA | Thyroid carcinoma | n=500, TCGA |
| UCEC | Uterine corpus endometrial carcinoma | n=167, TCGA |
| UCS | Uterine carcinosarcoma | n=111, TCGA |
| UVM | Uveal melanoma | n=79, TCGA |
| WT | Wilms tumors | n=130, TARGET |

**Figure S1**

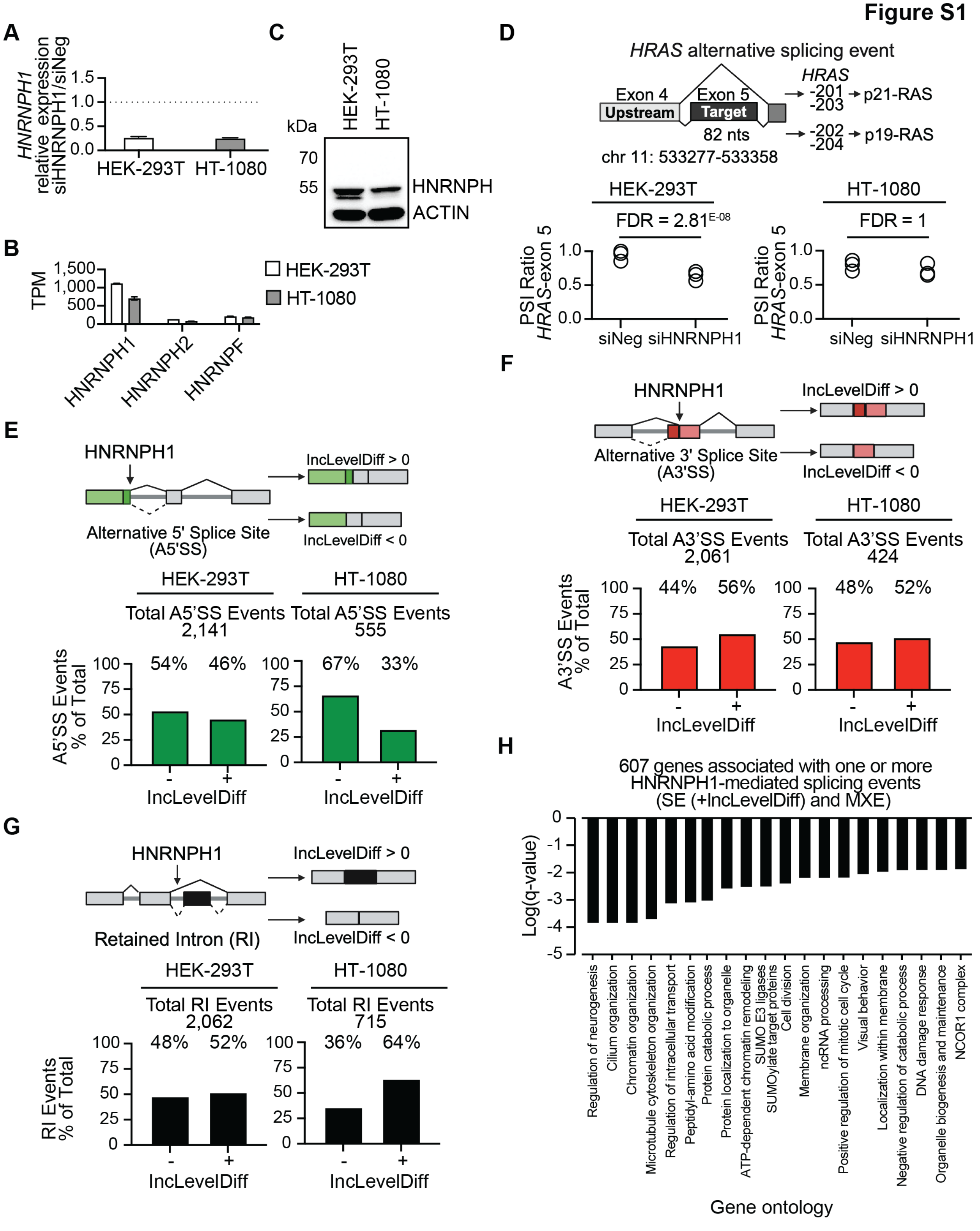


**Supplementary Figure S1. HNRNPH1's transcriptome-wide regulation of alternative splicing** (**A**) PCR-based quantification of *HNRNPH1* expression in siHNRNPH1-transfected HEK-

293T or HT-1080 cells normalized to siNeg-transfected cells. *HNRNPH1* expression normalized to *NACA*. (**B**) RNA-seq read counts for *HNRNPH1* and *HNRNPH2* in HEK-293T and HT-1080 cells. (**C**) Immunoblot analysis of whole cell lysates prepared from the indicated cell lines and probed using an antibody that due to the homology of HNRNPH1 and HNRNPH2 cannot distinguish between these proteins and an antibody against ACTIN as a loading control. Data shown is representative of three replicates. (**D**) A schematic of the *HRAS*-exon 5 splicing event that defines expression of transcript variants encoding the p21 and p19-HRAS isoforms and the rMATS analysis of the inclusion of the *HRAS*-exon 5 in HEK-293T and HT-1080 cells treated as indicated. (**E**) A schematic summarizing IncLevelDiff used to define HNRNPH1-dependent splicing events that make use of alternative 5′ splice sites and the percent distribution of IncLevelDiff values per cell line. (**F**) A schematic summarizing IncLevelDiff used to define HNRNPH1-dependent splicing events that make use of alternative 3′ splice sites and the percent distribution of IncLevelDiff values per cell line. (**G**) A schematic summarizing IncLevelDiff used to define HNRNPH1-dependent splicing events involving retained intron (RI) events and the percent distribution of IncLevelDiff values per cell line. (**H**) Gene ontology (GO) analysis of the 607 genes associated with one or more HNRNPH1-mediated splicing events (SE event, positive IncLevelDiff) and MXE events. Enrichment analysis performed using Metascape.

Figure S2

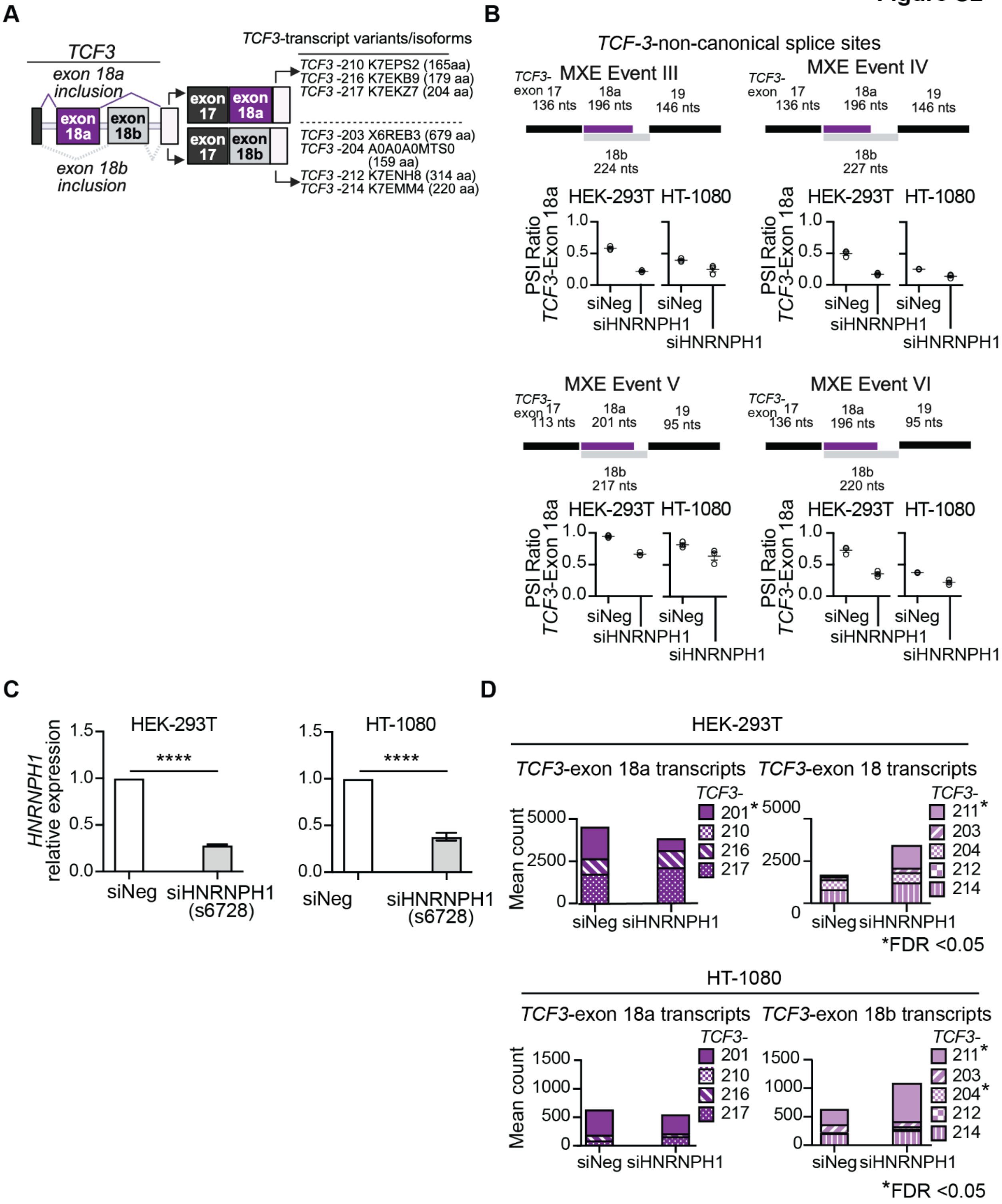


**Supplementary Figure S2: HNRNPH1's regulation of a *TCF3* mutually exclusive splicing event** (**A**) Schematic of *TCF3*-exon 18a and *TCF3*-exon 18b containing less abundant transcript variants and their encoded protein isoforms. (**B**) rMATS outputs for the *TCF3*-exon 18a and -exon 18b MXE splicing events that use non-canonical splice sites in the indicated cell lines transfected with the indicated siRNAs (three experimental replicas (open circles) and mean and SEM (lines). (**C**) qRT-PCR analysis of *HNRNPH1* expression in control (siNeg) and siHNRNPH1(s6728)-transfected HEK-293T and HT-1080 cells (unpaired t test with Welch's correction, P value **** <0.0001). (**D**) Quantification of *TCF3*-exon 18a or -exon 18b containing transcript variants expressed in control (siNeg) or HNRNPH1-depleted (siHNRNPH1) HEK-293T or HT-1080 cells (EBSeq). * Indicates FDR <0.05.

**Figure S3**

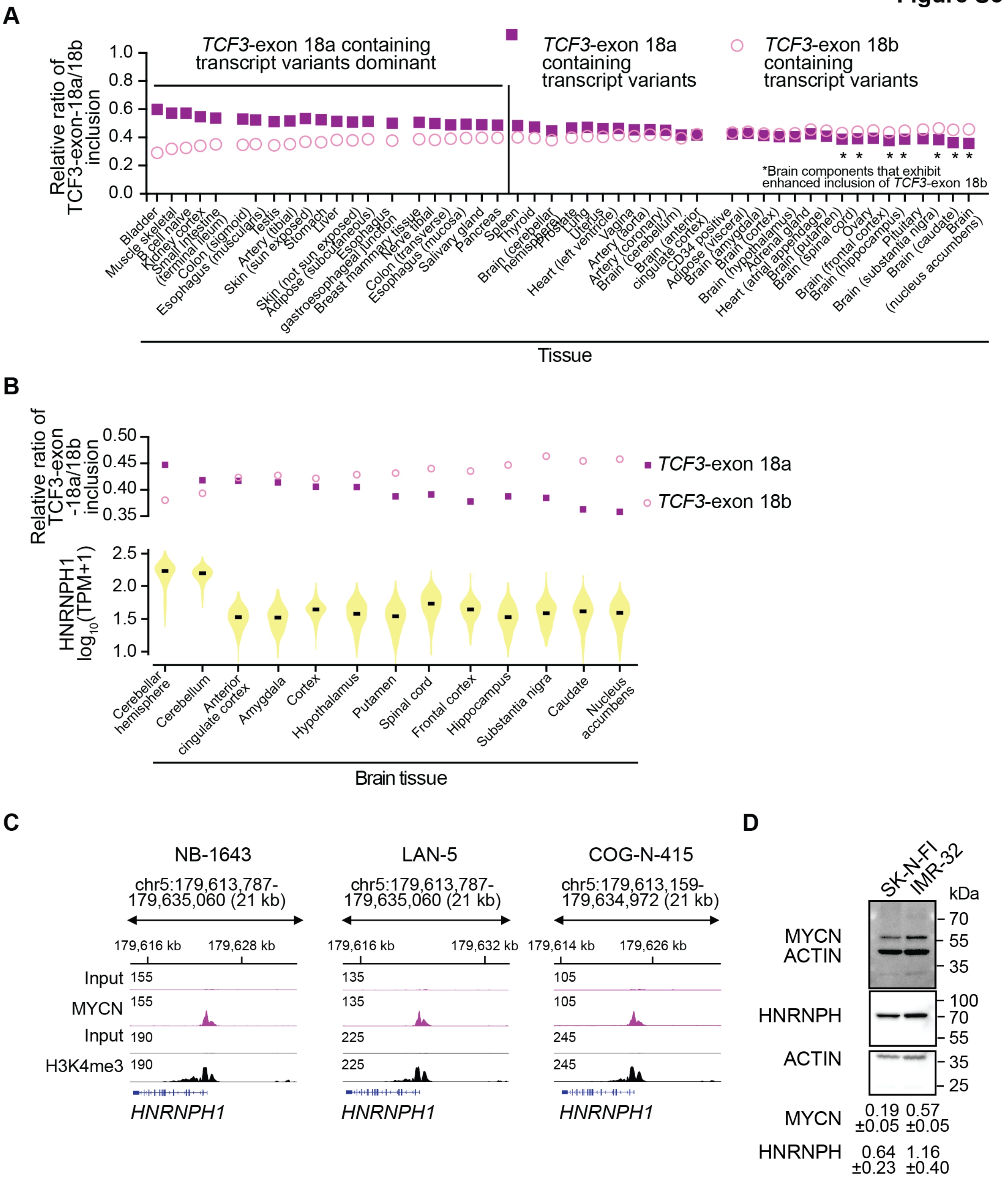


**Supplementary Figure S3: The differential expression of *TCF3* transcript variants in neuroblastoma** (**A**) Ranked relative ratios of *TCF3*-exon 18a and *TCF3*-exon 18b usage in the indicated tissues. Data extracted from MAJIQlopedia. (**B**) Ranked relative ratios of *TCF3*-exon 18a and *TCF3*-exon 18b usage and *HNRNPH1* expression in the indicated brain tissues (black bars indicate the median). Data extracted from MAJIQlopedia and GTEx portal, respectively (**C**) The *HNRNPH1* locus of NB-143, LAN-5 and COG-N-145 NB cells showing MYCN binding and H3K4me3 deposition (**D**) Immunoblot analysis of whole cell lysates extracted from the indicated NB cell lines probed using antibodies against MYCN and HNRNPH. Relative intensities (normalized to ACTIN) are shown on the bottom of each lane).

**Figure S4**

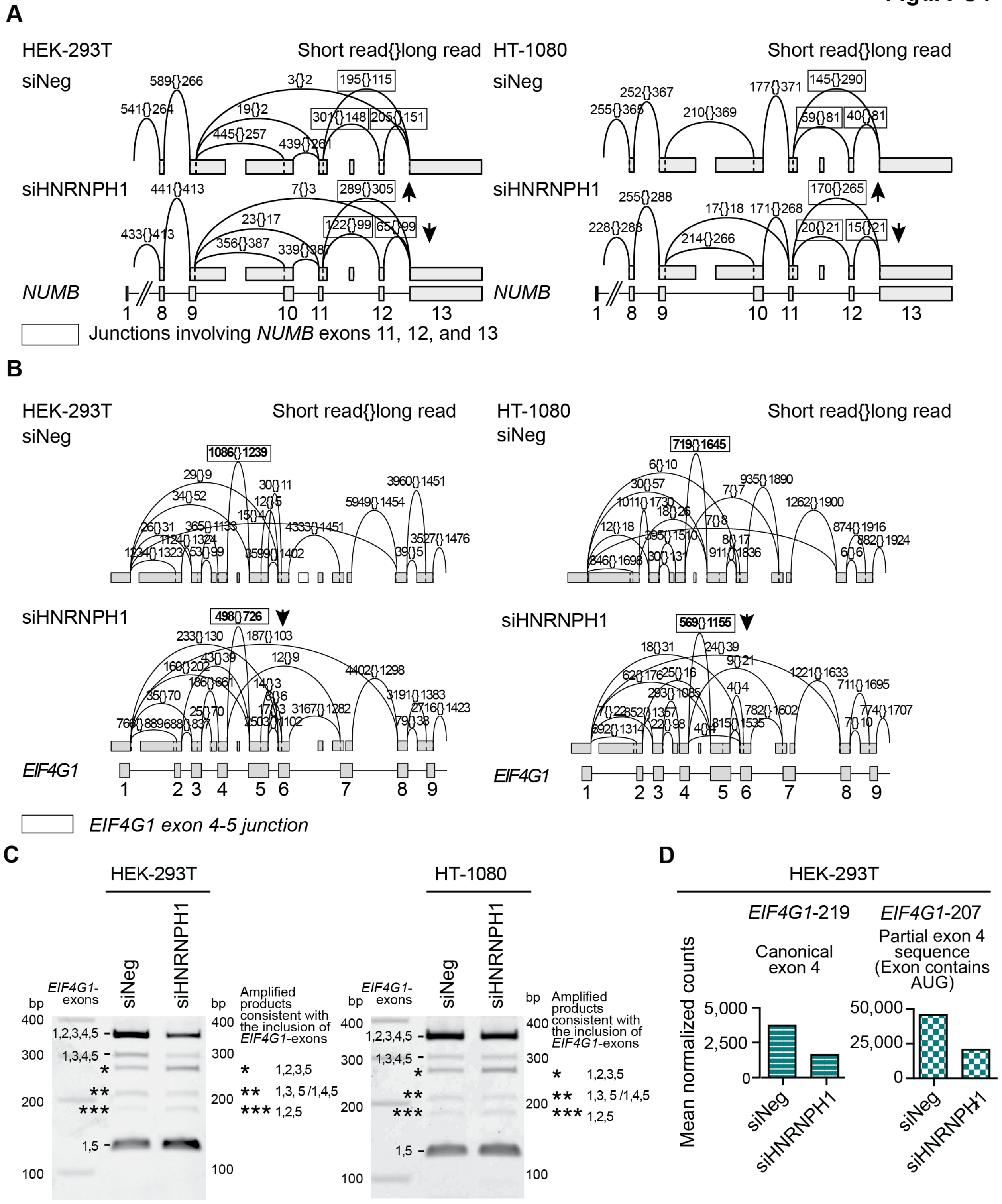


**Supplementary Figure S4: HNRNPH1 regulates the inclusion of *NUMB* exon 12 and usage of the *EIF4G1* exon 4-5 junction** (**A**) Visualization and quantification of splicing events focused on the 3′ *NUMB* exons (exons 9-13) expressed in HEK-293T and HT-1080 cells. The number on the left of each {} shows the median read count from the short-read RNA-seq data and the number on the right, the sum of full-length counts from the long-read data. The black rectangles indicate junctions involving NUMB exons 11, 12, and 13, and the arrows indicate the direction of change following the silencing of *HNRNPH1*. Data assessed using the VOILA visualization tool and quantified using the HET (heterogen) quantifier within MAJIQ (6). (**B)** Visualization and quantification of splicing events involving the first 9 exons of *EIF4G1* expressed in HEK-293T cells. The number on the left of each {} shows the median read count from the short-read RNA-seq data and the number on the right the sum of full-length counts from the long-read data. The black rectangles indicate the *EIF4G1* exon 4-5 junctions and arrows the direction of change following the silencing of *HNRNPH1*. Data assessed using the VOILA visualization tool and quantified using the HET quantifier within MAJIQ. (**C**) Representative longer exposures of the PCR-based splice assay designed to detect the usage of *EIF4G1* exons 1 – 5 in HEK-293T or HT-1080 cell lines transfected as indicated. The *, **, and *** specify amplified products predicted to consist of the indicated *EIF4G1* exons. (**D**) The mean normalized counts of the indicated *EIF4G1* transcript variants in control (siNeg) and HNRNPH1-depleted (siHNRNPH1) HEK-293T (EBSeq).

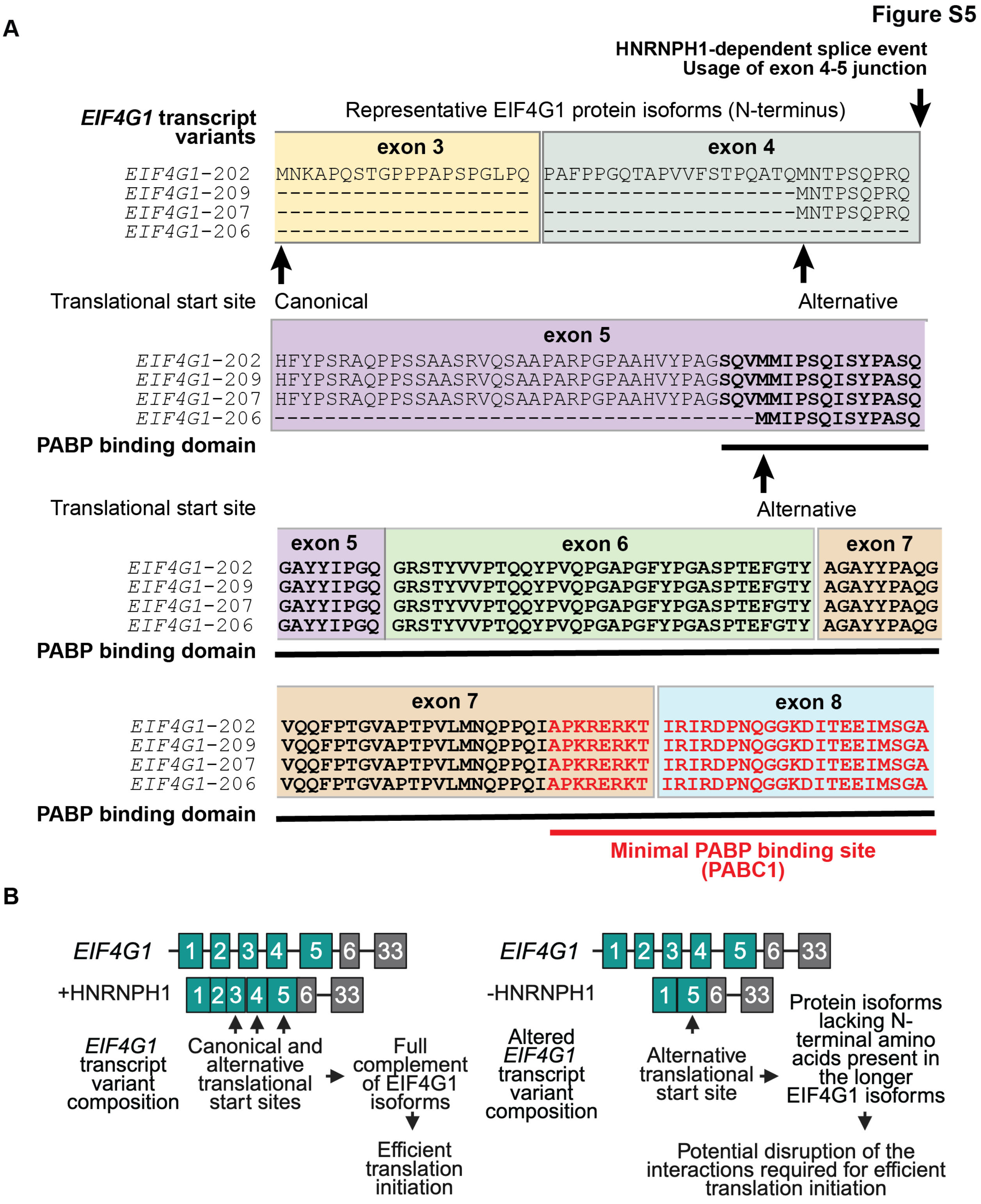


**Supplementary Figure S5: The depletion of HNRNPH1 reduces *EIF4G1* transcript variant composition** (**A**) The amino acid sequences of EIF4G1 isoform N-terminus regions encoded by the indicated transcript variants organized relative to the full-length EIF4G1 protein. The indicated exon annotations refer to the full-length *EIF4G1*-202 transcript variant. Information shown gathered from Ensembl gene ID ENSG00000114867 and UniProt entry Q04637. (**B**) A model illustrating HNRNPH1's contribution to the maintenance of EIF4G1 isoform composition through its regulation of the exon4/exon 5 splice event and the concept that HNRNPH1-depletion will alter EIF4G1 isoform composition by promoting the expression of isoforms lacking the amino acids which form the N-terminus of longer EIF4G1 isoforms. We speculate that this shift in isoform composition could disrupt the interactions required for efficient translation initiation.

**Figure S6**

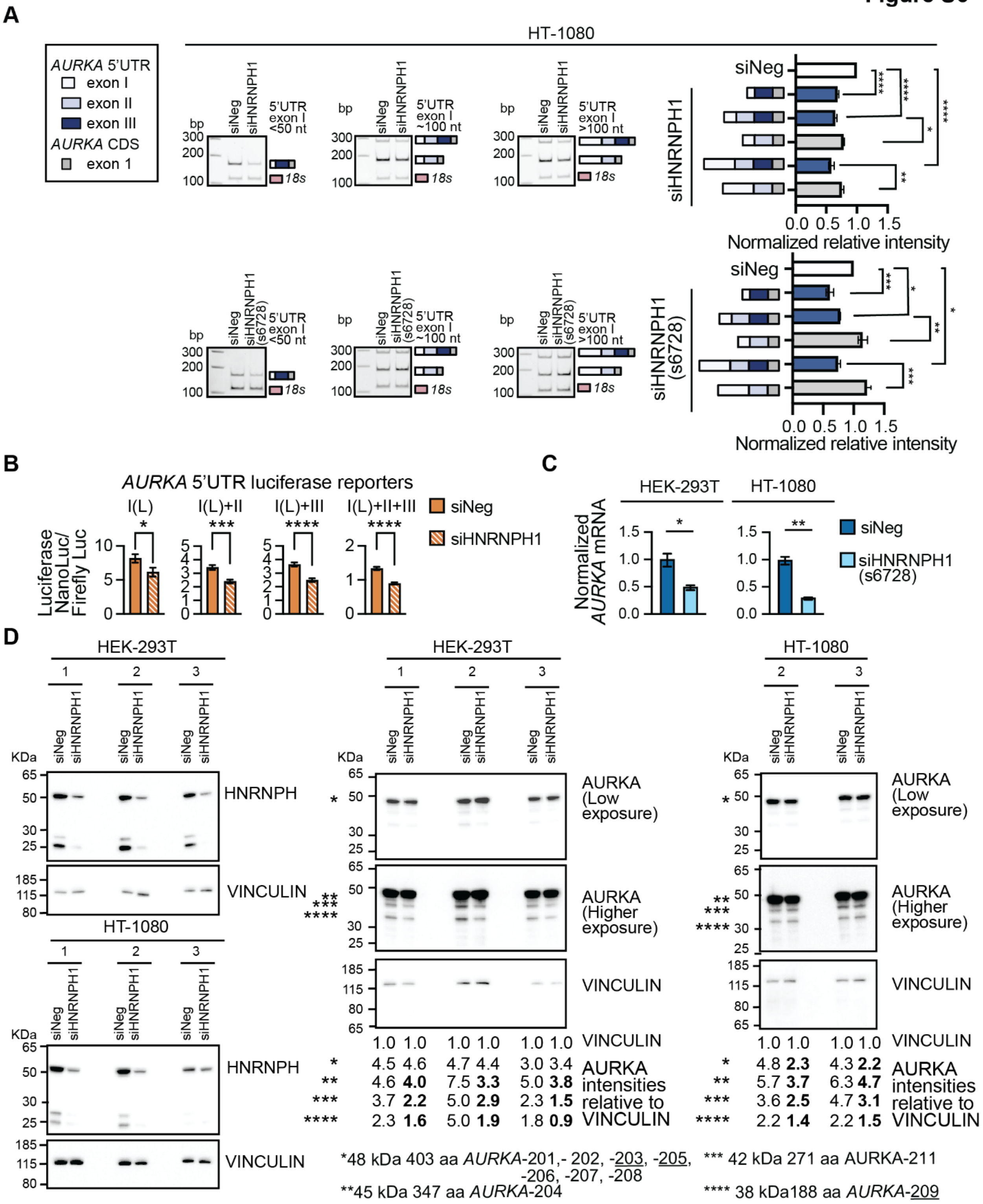


**Supplementary Figure S6: HNRNPH1's regulation of an *AURKA* 5'UTR skipped exon (SE) event and *AURKA* RNA stability.** (**A**) qPCR-based analysis of *AURKA* transcript variants that include *AURKA*-5'UTR-exon III in transfected HT-1080 cells (siNeg and siHNRNPH1: upper panels; siNeg and siHNRNPH1/s6278: lower panels), and quantification of the amplified products normalized to the intensity of the 18S amplified product (mean±SEM of three biological replicates). Statistical analysis: ordinary one-way ANOVA, P values *** <0.001, **** <0.0001. (**B**) Relative luciferase expression (NanoLuc/Firefly) 48 hours post-transfection of HEK-293T cells with the indicated reporter construct and the indicated siRNA (mean ± SEM n=6 per condition). Statistical analysis: unpaired t test with Welch's correction, P values * <0.05, *** <0.001, **** <0.0001. (**C**) qRT-PCR analysis of *AURKA* expression (*RPL27* normalized) following the silencing of *HNRNPH1*; unpaired t test with Welch's correction, P * <0.05, P ** <0.01. (**D**) The immunoblot analysis of whole cell lysates prepared from siNeg or siHNRNPH1(s6728)-transfected HEK-293T or HT-1080 cells probed using antibodies against the indicated protein. The images show the results of the three independent transfections per siRNA performed using HEK-293T cells and the two independent transfections per siRNA performed in HT-1080 cells. **Figure 7H** shows the results of the third transfection conducting using HT-1080 cells. The asterisks indicate the detected AURKA protein isoform and the values indicate the VINCULIN normalized quantification of each isoform.
